# metaJAM: a Nextflow integrated metagenomic workflow for sedimentary ancient DNA

**DOI:** 10.64898/2026.05.05.722689

**Authors:** Ernst Johnson, Chenyu Jin, Benjamin Guinet, Jamie Alumbaugh, Nathan Martin

**Affiliations:** Centre for Palaeogenetics, Stockholm, Sweden; Department of Geological Sciences, Stockholm University, Sweden; Department of Zoology, Stockholm University, Sweden, Sweden; Department of Bioinformatics and Genetics, Swedish Museum of Natural History, Sweden; Helmholtz Institute for One Health, Helmholtz Centre for Infection Research, Greifswald, Germany; Department of Microbiology, Immunology and Transplantation, Rega Institute, KU Leuven, Leuven, Belgium; Department of Archaeology and Classical Studies, Stockholm University, Sweden

**Keywords:** Metagenomic, ancient DNA, workflow, environmental DNA

## Abstract

The application of metagenomics in ancient DNA (aDNA) research is rapidly expanding, driven in particular by advances in sedimentary aDNA research and sequencing technologies. Although many ancient DNA studies rely on broadly similar bioinformatic strategies, there is still no single standardized, widely adopted workflow. These differences can directly affect how efficiently past biodiversity can be reconstructed and authenticated from the various archives analyzed using ancient metagenomic approaches.

Although a few pipelines tackle the processing of ancient DNA data from shotgun sequencing, the ones applied to metagenomic datasets are scarce and often resource-intensive or challenging to install, update, or extend with new tools and parameters. metaJAM, a scalable and user-friendly pipeline, is presented here to specifically address the challenges of metagenomic aDNA analyses of eukaryotes. The pipeline has been designed in Nextflow to ensure continuous development and can be used on different high-performance computing (HPC) clusters. metaJAM integrates all key steps required for ancient DNA metagenomic analyses, from raw sequencing data pre-processing to microbial filtering, taxonomic assignment via competitive iterative mapping against Bowtie 2 reference indexes and reassignment using lowest common ancestor (LCA) inference. Validation and authentication are performed using the post-LCA toolkit bamdam together with alignment to an exhaustive reference database using MMseqs2. It allows users to choose among alternative tools and generates a series of plots to support data visualization and taxon authentication.

metaJAM differs from existing pipelines through its implementation of rigorous filtering of microbial-like reads by Kraken 2 classification and masking microbial-like regions, iterative or parallel Bowtie 2 mapping, validation of the detected taxa and integration of up-to-date tools for ancient metagenomic analysis, along with diagnostic plots that help users assess the reliability of taxonomic assignments and visualize their data. It complies well with limited computational resources, customised databases for taxonomical groups, and provides an accessible workflow to support the investigation of metagenomic ancient DNA datasets. Its applications span a range of contexts, from ecosystem reconstructions in environmental aDNA archives such as sediments, to metagenomic studies on archaeological artefacts and even taxonomic identification of undiagnosed biological materials.

## Introduction

Building on pioneering studies of genetic material preserved in ancient environmental samples (Coolen and Overmann, 1998; Poinar et al., 1998), metagenomic analyses of ancient environmental DNA (eDNA) have become a widely used and effective approach for reconstructing past ecosystem composition and enabling genetic investigations of ancient communities (Murchie et al., 2021; Orlando et al., 2021; Thomas et al., 2012). Processing ancient DNA sequencing data requires accounting for the distinctive properties of ancient DNA molecules. Retrieved sequences are typically highly fragmented and degraded due to the taphonomic processes acting on DNA after an organism’s death (Briggs, 2020; Dabney et al., 2013; Gilbert et al., 2007; Lindahl, 1993; Willerslev and Cooper, 2004), and endogenous DNA is often present in low amounts, increasing the risk of contamination from exogenous sources in the laboratory or surrounding environment. Consequently, read lengths are generally short, often below ca. 150 base pairs (bp) (Meyer et al., 2012; Poinar et al., 2006; Renaud et al., 2019), and they commonly exhibit characteristic deamination damage patterns. In particular, cytosine deamination converts cytosine to uracil, which is then read as thymine during library amplification, resulting in a characteristic excess of cytosine-to-thymine (C-to-T) substitutions at the ends of DNA fragments (Gilbert et al., 2007). Although these specific characteristics can help authenticate the ancient origin of taxa detected in metagenomic studies, they also necessitate dedicated bioinformatic processing steps and tailored workflows (Orlando et al., 2021).

Ancient metagenomic analysis generally involves a series of data-processing steps intended to improve data quality and support reliable biological interpretation. The first general step is the cleaning of raw sequencing data to remove technical artefacts and biases introduced during library preparation and sequencing by removing low-complexity and PCR-derived duplicate reads, using tools such as SGA v0.10.15 (Simpson and Durbin, 2012) or PRINSEQ (Schmieder and Edwards, 2011).

Overlapping read pairs are also merged during the pre-processing treatments of the sequencing reads (Kircher, 2012). A wide range of software is available for these preprocessing steps (Orlando et al., 2021), and the specific choices often vary depending on author preferences. The processed reads are then taxonomically assigned to their most likely taxonomic origins, most commonly by alignment tools using a competitive mapping strategy such as BWA or Bowtie 2, the latter being preferred for metagenomic studies due to its speed and allowance of multimapping sequences (Oliva et al., 2021; Poullet and Orlando, 2020). This assignment step is critical because the fragmented and damaged nature of ancient DNA makes accurate mapping to specific taxa challenging (Dolenz et al., 2024; Yates et al., 2021) as one must account for mismatches and gaps, as well as biases introduced by reference choice and spurious mapping. All of which can influence both taxonomic identification and the detection of authentic ancient DNA damage signatures (Dolenz et al., 2024; Oliva et al., 2021).

Misassignment, whether driven by contamination or issues in the reference database, is particularly difficult to address in metagenomic datasets. Using more exhaustive reference databases can help mitigate this problem (Key et al., 2017) and reduce the detection of “oasis” taxa (Cribdon et al., 2020). However, it is important to note that reference collections are often biased toward organisms of particular interest, for example, those relevant to agriculture or medicine (Michalcová et al., 2011; Troudet et al., 2017), and they may also contain errors due to limited curation (International Society for Biocuration, 2018; Orlando et al., 2021). To mitigate this, an additional lowest common ancestor (LCA) step, using software such as ngsLCA (Wang et al., 2022), is generally applied to resolve taxonomic identities across ranks. After sequences assigned to taxa of interest are identified, some metagenomic ancient DNA studies further perform an alignment-based classification against a more exhaustive database (Huson et al., 2016) using BLAST (Altschul et al., 1990) or the faster recent alternative MMseqs2 (Steinegger and Söding, 2017). Running these algorithms on a reduced, refined dataset is less time and resource-intensive.

Finally, to further authenticate ancient taxonomic assignments and to reduce false positive taxonomic assignments, studies commonly examine damage patterns and read-length distributions (Armbrecht et al., 2021; Jónsson et al., 2013; Skoglund et al., 2012) and apply minimum read-count or relative-abundance thresholds to support a taxon’s presence (Pedersen et al., 2016; Slon et al., 2017; Stahlschmidt et al., 2019). However, these filters can remain too permissive, as false positives may also arise from low-complexity and contaminant microbial-like regions incorporated within reference eukaryote genomes, motivating upstream masking of problematic regions in reference databases (Laurence et al., 2014; Mann et al., 2023; Merchant et al., 2014; Oskolkov et al., 2025). More recently, the tool bamdam also helped to address this issue by providing a post-mapping, post-LCA authentication toolkit (De Sanctis et al., 2025). Several pipelines are available for processing ancient DNA shotgun sequencing data, but many of the earlier workflows such as EAGER (Peltzer et al., 2016) or PALEOMIX (Schubert et al., 2014) were not designed for metagenomic applications and are not straightforward to extend when incorporating new tools or modifying parameters. More recently, a small number of pipelines have been developed to support metagenomic ancient DNA analyses, including nf-core/eager (Yates et al., 2021), HAYSTAC (Dimopoulos et al., 2022), aMeta (Pochon et al., 2023), Quicksand (Szymanski et al., 2025), and Holi (Kjær et al., 2022; Pedersen et al., 2016; Vogel et al., 2026). nf-core/eager integrates HOPS (Hübler et al., 2019), which is primarily intended for automated bacterial screening and can be challenging to map against large reference databases. aMeta, while substantially faster and less resource-demanding thanks to its KrakenUniq-based screening, similarly prioritizes microbial targets in its scoring system and is currently less suited to eukaryotic identification. In contrast, Holi is tailored towards eukaryotes, but typically requires considerable computational resources and large reference databases that may be difficult for many research groups to access due to limited computational resources, and its execution can be complex. Quicksand is optimized for mammalian mitochondrial DNA in metagenomic eDNA datasets and is therefore not intended to cover other organismal groups, such as Viridiplantae, which are often essential for ecosystem reconstruction.

To address the challenges of metagenomic analyses in ancient DNA, we present here metaJAM, which is a simplified, scalable, reproducible and user-friendly Nextflow pipeline (Di Tommaso et al., 2017) specifically designed for eukaryote detection. The integration of up-to-date tools for ancient metagenomic analysis and database creation, paired with diagnostic plots, aims to help the user to assess the reliability of taxonomic assignments and visualize their data. metaJAM is designed to run efficiently on high-performance computing (HPC) systems, with customizable inputs that can be refined to optimize performance and runtime. It also supports parallel execution, enabling the simultaneous analysis of multiple ancient eDNA samples.

## Materials and Methods

### Workflow

The pre-processing sections of metaJAM consist first of the removal of sequencing adapters using fastp (Chen et al., 2018), including adapter trimming, poly-X tail trimming (trim_poly_x), and merging of overlapping forward and reverse reads, outputing merged and unmerged reads together for downstream analysis.The fastp parameters can be customized (e.g., overlap_len_require or the minimum read length), while the defaults commonly used in ancient DNA studies are already implemented in the workflow. In a second step, low-complexity reads and PCR duplicates are filtered using either PRINSEQ or SGA (Schmieder and Edwards, 2011; Simpson and Durbin, 2012). Similarly, aDNA-suited default settings are provided, while key parameters such as the DUST threshold for complexity filtering and minimum read length remain user-adjustable.

Microbial contaminants are then removed from the processed reads using a k-mer-based classification approach with Kraken 2 v2.17.1 (Wood et al., 2019) against an exhaustive microbial reference database. To maximize reproducibility and coverage of microbial genomes, use of the GTDB resource (Parks et al., 2026) is recommended. metaJAM can also hard-mask microbial contamination found in reference fasta files through a module based on the GENEX workflow described in Oskolkov et al. (2025). This module creates bed files containing the coordinates of microbial-like regions found within the references, which can then be used to mask out potential contamination either a) in the reference fasta files prior to index building out of metajam and then supply the mapping index to metajam, or b) in metajam to mask bam files produced after mapping. Kraken 2 databases built with the default k-mer size (35 bp) do not classify reads shorter than 35 bp and tend to show reduced classification rates for short reads, potentially missing microbial (or microbial-like) sequences. Therefore, we recommend the use of both the Kraken 2 and GENEX methods of masking, as the latter could exclude reads of any length as long as it locates in microbial-like reference regions.

Classified reads are then aligned to the user’s reference databases using the aligner Bowtie 2 allowing outputing the reads best mapped to multiple genomic locations (Langmead and Salzberg, 2012). The workflow offers users the choice of either iterative or parallel mapping. In the iterative, funnel-like strategy, one or more databases are processed sequentially, where reads that remain unmapped against the first database are passed to the next, and so on. Alternatively, in the parallel mapping strategy, all pre-processed reads are mapped to each database simultaneously and equally. This approach is suitable when no a priori knowledge exists regarding which database contains higher-quality genomes, best represents local biodiversity, or is of greater research interest compared with the other databases used. The mapping results against all databases are merged per sample and sorted using SAMtools v1.20 (Li et al., 2009).

After merging, each read is assigned to the lowest supported taxonomic node using the LCA software ngsLCA (Wang et al., 2022). This step relies on a custom acc2taxid file containing the contig and their NCBI taxonomical ID of all reference databases used during mapping, together with the NCBI taxonomy names and nodes files (Schoch et al., 2020). Optionally, read alignments can be refined prior to LCA assignment using filterBAM (https://github.com/genomewalker/bam-filter). To assess authenticity and support taxa validation, the post-LCA authentication toolkit bamdam v0.4.3 (De Sanctis et al., 2025) can then be applied to ngsLCA-classified reads using either recommended or user-defined bamdam settings.

As an additional layer of evaluation, MMseqs2 v17.b804f search using the BlastN-like mode (Steinegger and Söding, 2017), can be used to query a subset of reads assigned to taxa of interest against a large reference database, preferably the NCBI nt database. Shotgun metagenomics can in principle recover enough genetic information to identify organisms down to the species or even individual level (Johnson et al., 2026). In practice, however, taxonomic classification of aDNA data is often limited to the genus level or above, as the short and degraded nature of ancient DNA fragments reduces confidence in species-level assignments (Heintzman et al., 2023). MMseqs2 can be run in metaJAM either for the 10 most abundant genera in each identified kingdom, or with an a priori query list of specific taxa of interest that the user wants to investigate. Bamdam and MMseqs2 taxonomic assignments are then compared in order to evaluate the confidence of final taxonomic rank identification. In the event that references for a genus of interest are not available in the NCBI nt-database, metaJAM will flag the problem and prompt users to explore other options, such as using BLAST against a more taxa specific NCBI Whole Genome Shotgun (WGS) database, to complete their investigation. At the end of the pipeline, users receive easy-to-read visual summaries that combine this feedback with taxonomic assignment evaluation. In addition to bamdam outputs, optional visualization modules generate summary plots of detected taxa and key processing metrics using custom Python and R scripts and a color-blind-friendly palette (Ichihara et al., 2008). Heatmaps and bubble plots help users inspect damage patterns, assess assignment quality, and judge the authenticity and reliability of reported taxa. Finally, up-to-date versions of the required tools are used to develop metaJAM and the dependencies are deployed using conda-based environments to promote stability and reproducibility.

### Usage

metaJAM is designed to run on HPC clusters and requires Nextflow dependencies together with conda (https://conda.io). Once these prerequisites are in place, Nextflow manages the pipeline’s software dependencies, providing users with a straightforward and reproducible installation set up. It also helps users to optimize compute allocation (CPU/memory/time) with automatic retry mechanisms that dynamically scale resource requirements up to the limits of the cluster.

To run metaJAM, users specify the required processes, input files, and parameters to be executed. The pipeline can then be launched with a single command, and each process can also be executed independently. Raw sequencing data can be provided as gzipped FASTQ files to run the complete workflow; alternatively, metaJAM accepts intermediate inputs from any step of the pipeline in gzipped FASTQ or BAM format. Reference databases are supplied in indexed Bowtie 2 formats, and a metadata sheet in TXT format is required to enable plot generation and visualization.

### Benchmark dataset

To illustrate the use of metaJAM in metagenomic ancient DNA research, we evaluated the pipeline on a benchmark dataset containing metagenomic samples from Courtin et al. (2022), where the authors studied permafrost sediments from Batagay megaslump in East Siberia. Three samples representing glacial ecosystems, B17-D6_MIS3, B17-D7_MIS3, and B17-D10_MIS3, respectively dated to 27 ky BP, 23 ky BP and 50 ky BP, were chosen based on the numerous Viridiplantae and Mammalian reads previously identified in those layers using a classifier approach based on Kraken 2 (Wood et al., 2019). The raw sequencing data were retrieved from the European Nucleotide Archive Project Number PRJEB43506, and only the deep sequenced samples (Run accessions ERR9690362, ERR9690364, ERR9690365) were analyzed through metaJAM.

Different parameters were set in the config file from metaJAM nextflow workflow. First, the paired-end read files were trimmed and merged using fastp v0.24 (Chen et al., 2018), allowing for 20 bp overlap and a minimum read length of 30 bp. Second, low-complexity reads with a high DUST score (> 4) and exact forward and reverse duplicates (option 14) were removed with PRINSEQ v0.20.4 (Schmieder and Edwards, 2011). After pre-processing, both the merged and unmerged reads were passed to Kraken 2 v2.1.2 using default parameters (Wood et al., 2019) to classify the reads against the Genome Taxonomy database (GTDB) release 226 (Parks et al., 2026). The reads were then mapped using Bowtie 2 v2.5.4 (Langmead and Salzberg, 2012), allowing for 1000 alignments per read (-k 1000) against different databases. To better evaluate the efficiency of the workflow towards different kinds of databases, complete genomes and nuclear dataset were added to the original mitochondrial and chloroplast RefSeq databases used in the last Kraken classification step in Courtin et al. (2022).

In total, the processed reads were mapped iteratively to four different databases using a funnel strategy, starting from the most complex database with the most potential matches to the sediment aDNA taxa composition, to the least complex database with the fewest likely matches. The first database used was PhyloNorway (Alsos et al., 2020) as provided in Wang et al. (2021), which was masked for microbial presence using the metaJAM module for the GENEX workflow (Oskolkov et al,. 2025). Then remaining unmapped reads were aligned against the plastid RefSeq created in 2025-05-23, followed by the mitochondrial RefSeq databases created in 2025-07-11 (Goldfarb et al., 2025), and finally by a custom database containing complete genomes for potential mammalian, fish and bird targets (Supplementary Table 1).

The mapped reads were taxonomically assigned using ngsLCA v0.9 (Wang et al., 2022) with the parameters -simscorelow 0.95 and -simscorehigh 1. Ancient DNA damage profiles, based on nucleotide base transitions of C to T and G to A, were calculated and visualised by bamdam v0.4.2 (De Sanctis et al., 2025). A slight local modification was applied to the bamdam krona python script, to visualize the damage per taxa in the krona plot as the average of the Damage +1 and Damage -1 values obtained from the bamdam compute step. The average DNA damage percentage was evaluated per sample, and taxa with less than 3.5 % damage were determined as not ancient. As a final validation, a random subset of up to 200 classified reads from each taxon identified in Courtin et al. (2022) alongside the most abundant detected per sample were blasted with MMseqs2 search (Steinegger and Söding, 2017) against the NCBI nt database (version 20240202) with the following settings: a maximum of 300 hits per sequence, a sequence identity threshold of 85 %, minimum read coverage of 95 %, maximum E-value of 1e-5, minimum bit score of 50, and a sensitivity of 7.5. A minimum of 10% difference of the bitscore between the two best hits of different genus was applied to evaluate the taxonomic assignment, applying a similar filter than Murchie et al. (2021) in the MEGAN analysis of BLAST results. The concordance between MMseqs2 and bamdam taxonomic assignments was evaluated using empirically defined thresholds and visualized accordingly. Additional MMseqs2 search analysis against a taxon-specific whole-genome shotgun (WGS) database was deemed necessary when more than 50% of the queried reads were classified as no hits. A taxon was considered confidently assigned at the genus level when at least 40% of reads with one or more hits shared the same genus-level assignment as that obtained with bamdam. Family-level assignment was considered supported when at least one third of reads with one or more hits matched the bamdam assignment at the family level. In such cases, additional investigation was recommended to refine genus-level identification. Taxa that did not meet any of these criteria were classified as likely false assignments, potentially reflecting database bias or contamination.

## Results

When the metajam workflow was run on three samples from Courtin et al. (2022), the runtime and memory usage remained below 60 GB, except for processes that required high-memory compute nodes. These computationally intensive steps included Kraken2 classification, Bowtie 2 mapping, and the MMseqs2 alignment step, primarily due to the large reference databases involved (Supplementary Fig. 1).

### Reads processing

Following adapter trimming and quality filtering with fastp, the shotgun libraries contained an average of 225,708,478 reads (SD ± 9,793,339). After removal of PCR duplicates and low-complexity reads using PRINSEQ, 157,321,093 reads remained on average (SD ± 4,800,049).

Classification against the Kraken 2 GTDB database indicated that 39.3% of reads were of bacterial origin, while 61,806,764 reads (SD ± 2,492,204) remained unclassified. In total, 12,337,339 (SD ± 945,790) unclassified reads mapped to the combined reference databases; most aligned to the PhyloNorway database (99.85%), with smaller fractions mapping to RefSeq (0.08%) and to the custom database of complete mammalian and fish genomes (0.07%). Subsequent post-mapping processing with ngsLCA and bamdam yielded an average of 8,136,673 assigned reads (SD ± 753,916) (Supplementary Fig. 2).

To authenticate each taxon of interest, the metaJAM workflow output the bamdam damage and Krona plots (Supplementary Fig. 3), which reported the proportion of reads with a C to T substitution at the 5’ end, as well as the DUST, complexity, and mean length metrics alongside decision-advice plots delivered by the pipeline based on the damage (Supplementary Fig. 4) and the MMseqs2 results (Supplementary Fig. 5).

### Viridiplantae composition

Most of the assigned taxa belonged to the Viridiplantae, with the assigned plant assemblage dominated by the families Poaceae and Asteraceae (Figs. 3A–B). At the genus level, Artemisia and Plantago were the most abundant taxa (Figs. 3D, 3F). No tree taxa were detected except for Salix, observed in samples B17-D6 and B17-D10 (Fig. 3D), which likely reflects shrub willow rather than tree cover according to the context of the samples (Supplementary Fig. 6D-E). Overall, these results are consistent with Courtin et al. (2022), which described a plant community characteristic of glacial steppe ecosystems. Comparison of taxa detected or not by Courtin et al. (2022) with those authenticated using metaJAM showed similar qualitative results (Fig. 2). The number of reads assigned per taxa were higher in the metaJAM strategy, principally due to the inclusion of nuclear skim genomes references in the microbially-masked PhyloNorway database beside organelle genomes and an updated version of the RefSeq plastid database. The use of these datasets resulted in the detection of some previously undetected genera, notably Salix, Rumex, and Dryas (Supplementary Fig. 6C, damage patterns). Conversely, the few taxa detected by Courtin et al. (2022) but not retrieved in metaJAM were rare and largely restricted to species-level assignments. This discrepancy is consistent with methodological differences, as the ngsLCA and bamdam post-mapping authentication steps used in metaJAM tend to yield more conservative assignments and often support higher taxonomic ranks than Kraken 2. The discrepancy in performance can be attributed to the higher accuracy and recall rate at species level of Bowtie 2 than Kraken 2 (default: confidence threshold=0), as recently shown by Gao et al. (2025) based on simulated metagenomic data for three species of rice.

**Figure 3.A.**
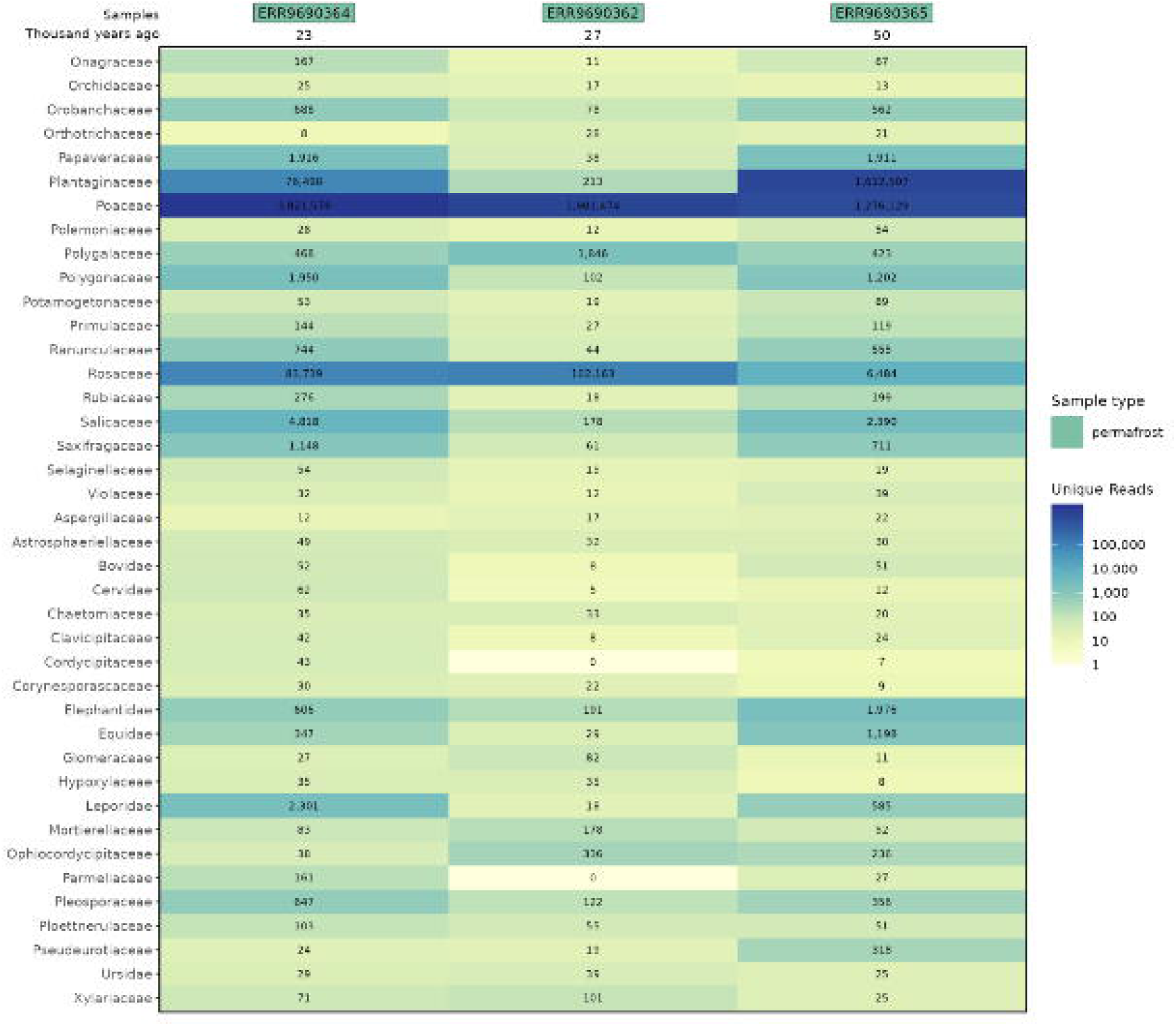
Heatmap representing the number of unique reads assigned per taxa at the family level after the bamdam step (1 to 50)

**Figure 3.B.**
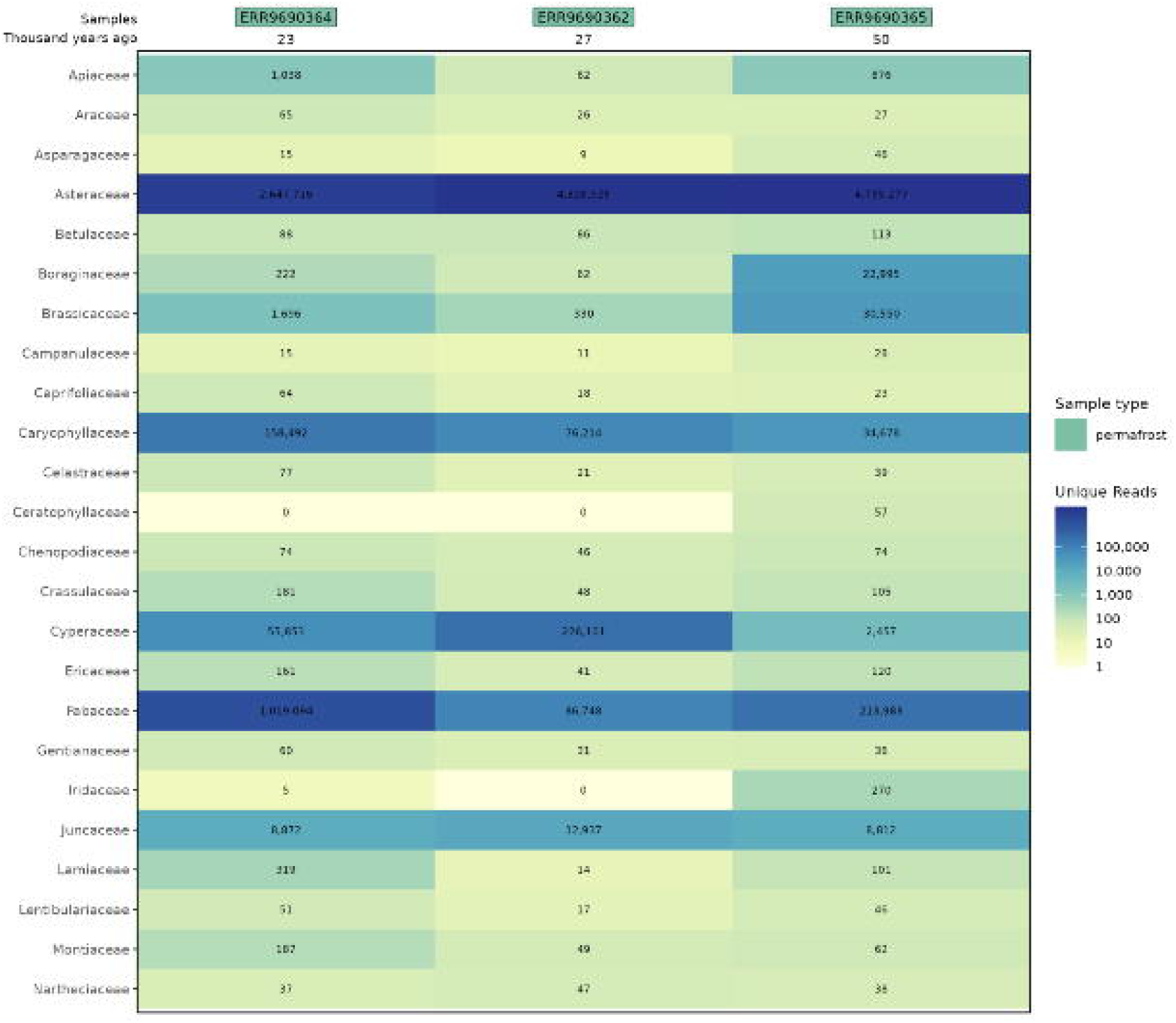
Heatmap representing the number of unique reads assigned per taxa at the family level after the bamdam step (51 to 77)

**Figure 3.C.**
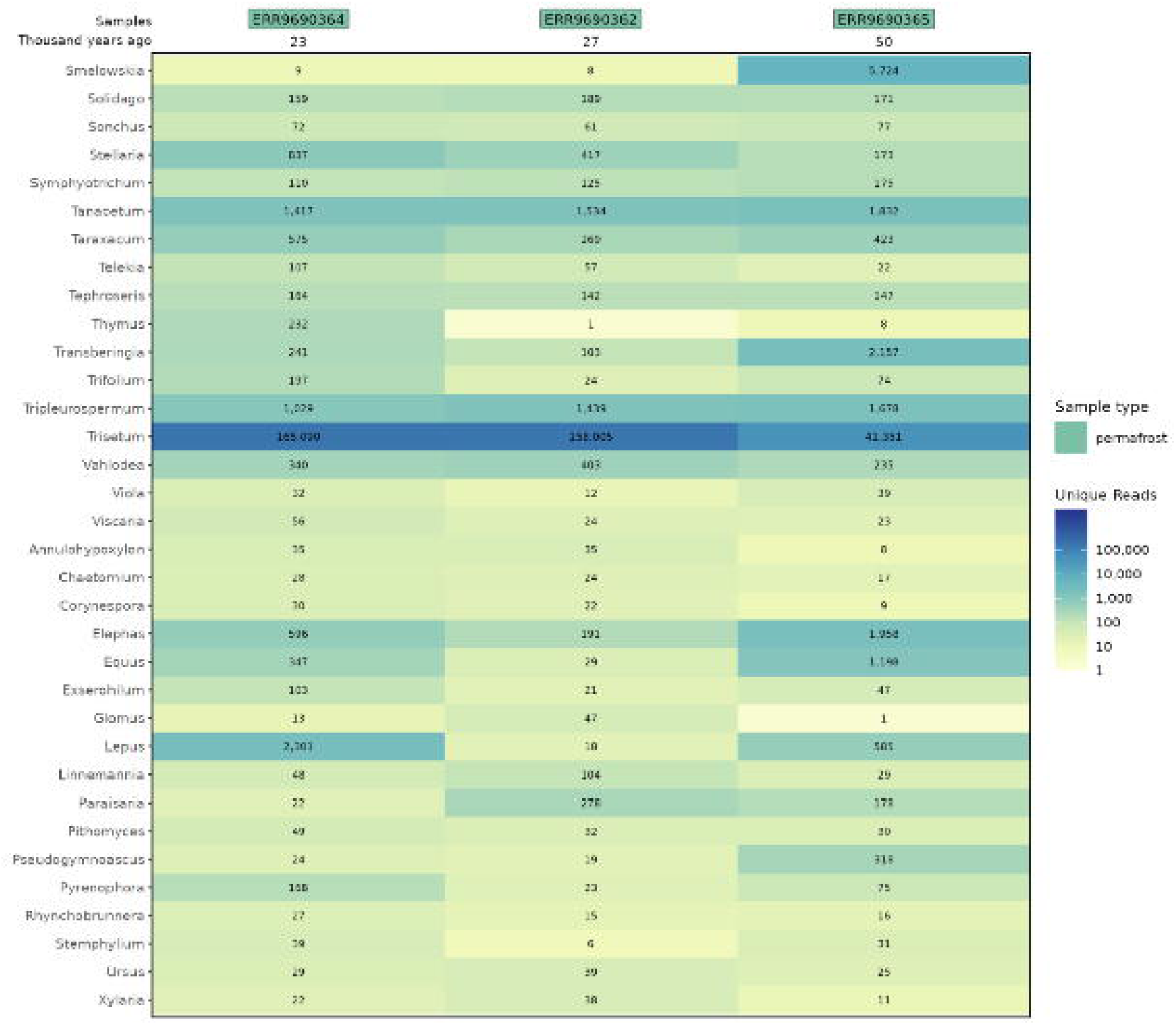
Heatmap representing the number of unique reads assigned per taxa at the genus level after the bamdam step (1 to 50)

**Figure 3.D.**
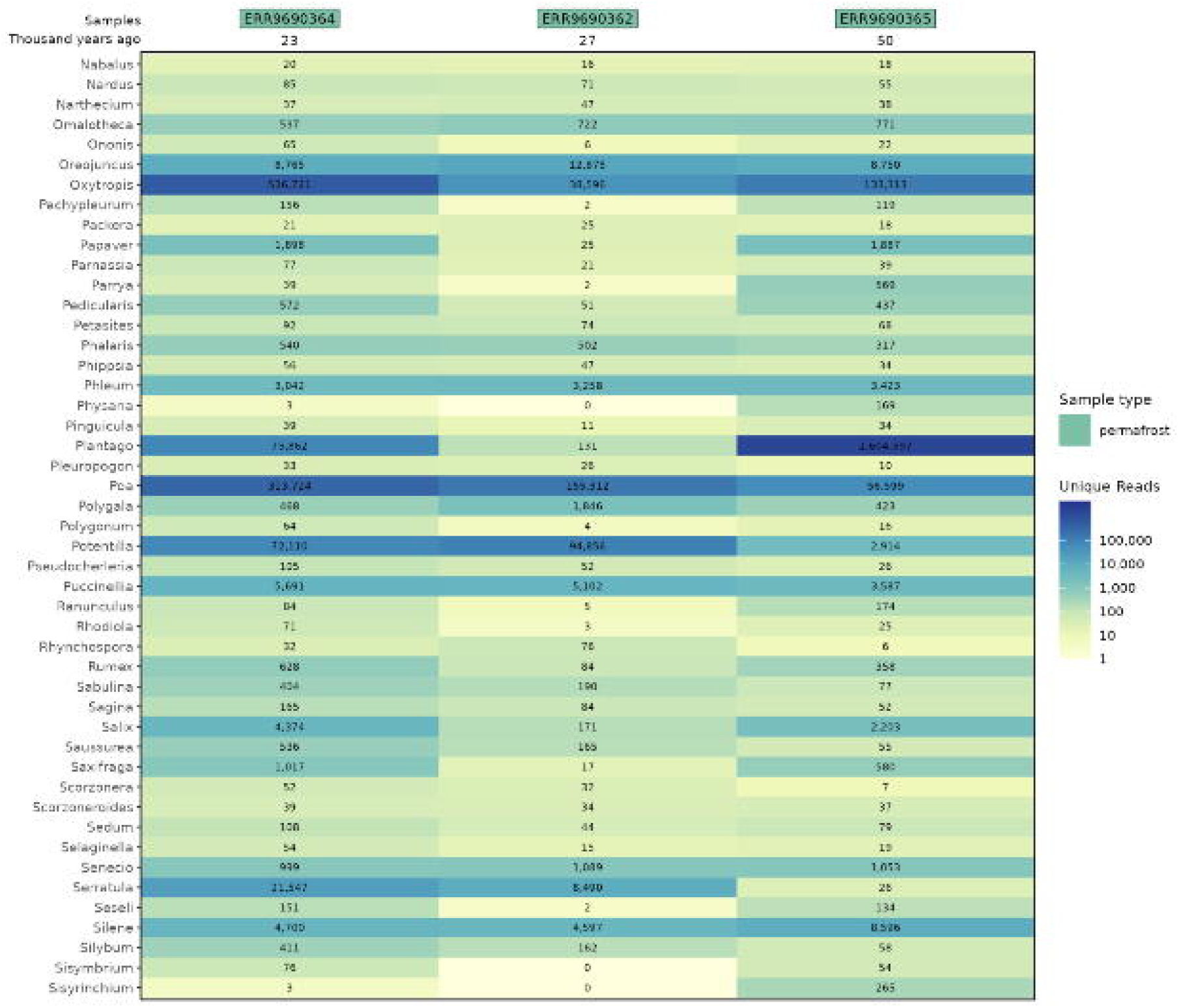
Heatmap representing the number of unique reads assigned per taxa at the genus level after the bamdam step (51 to 100)

**Figure 3.E.**
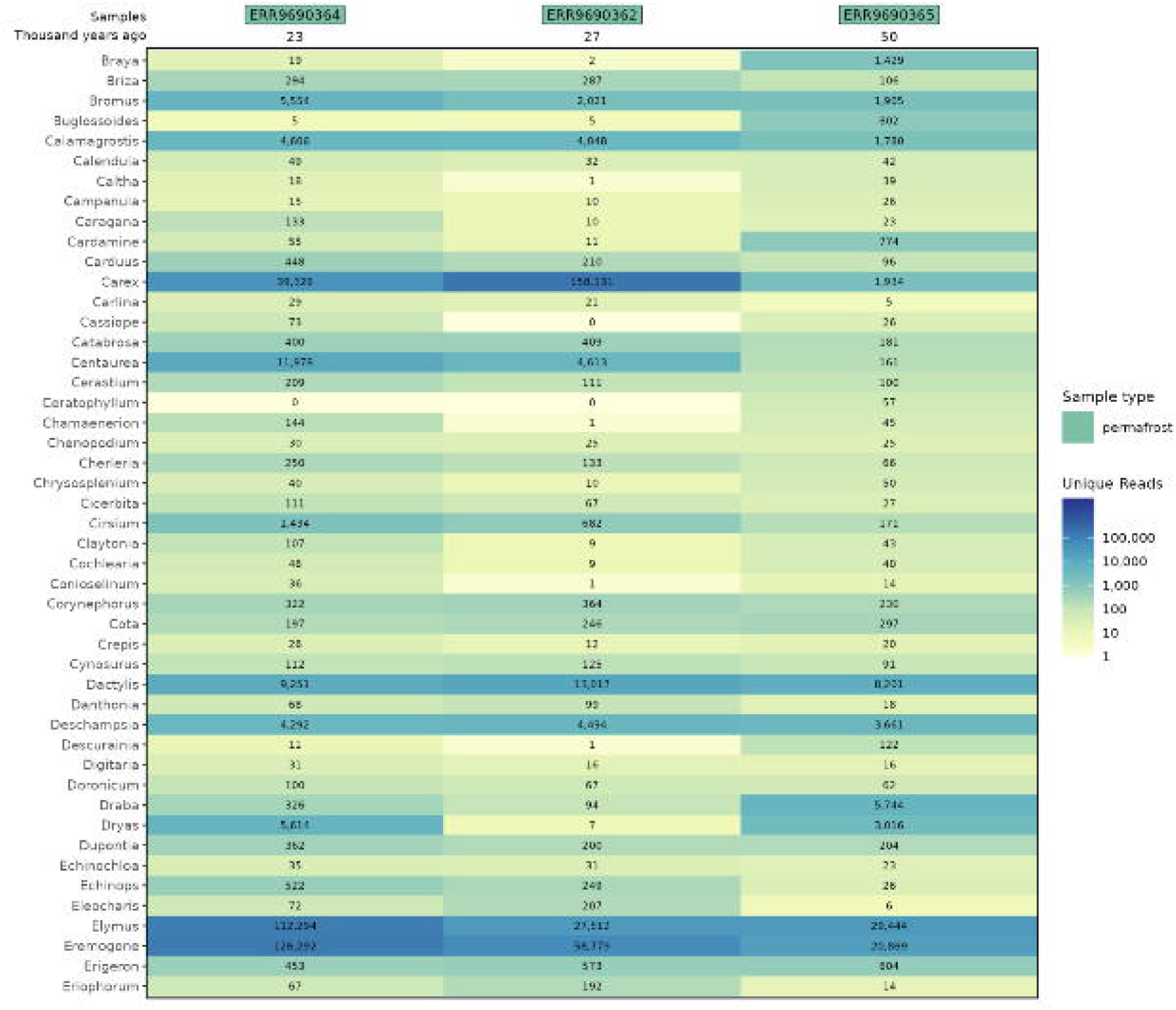
Heatmap representing the number of unique reads assigned per taxa at the genus level after the bamdam step (151 to 200)

**Figure 3.F.**
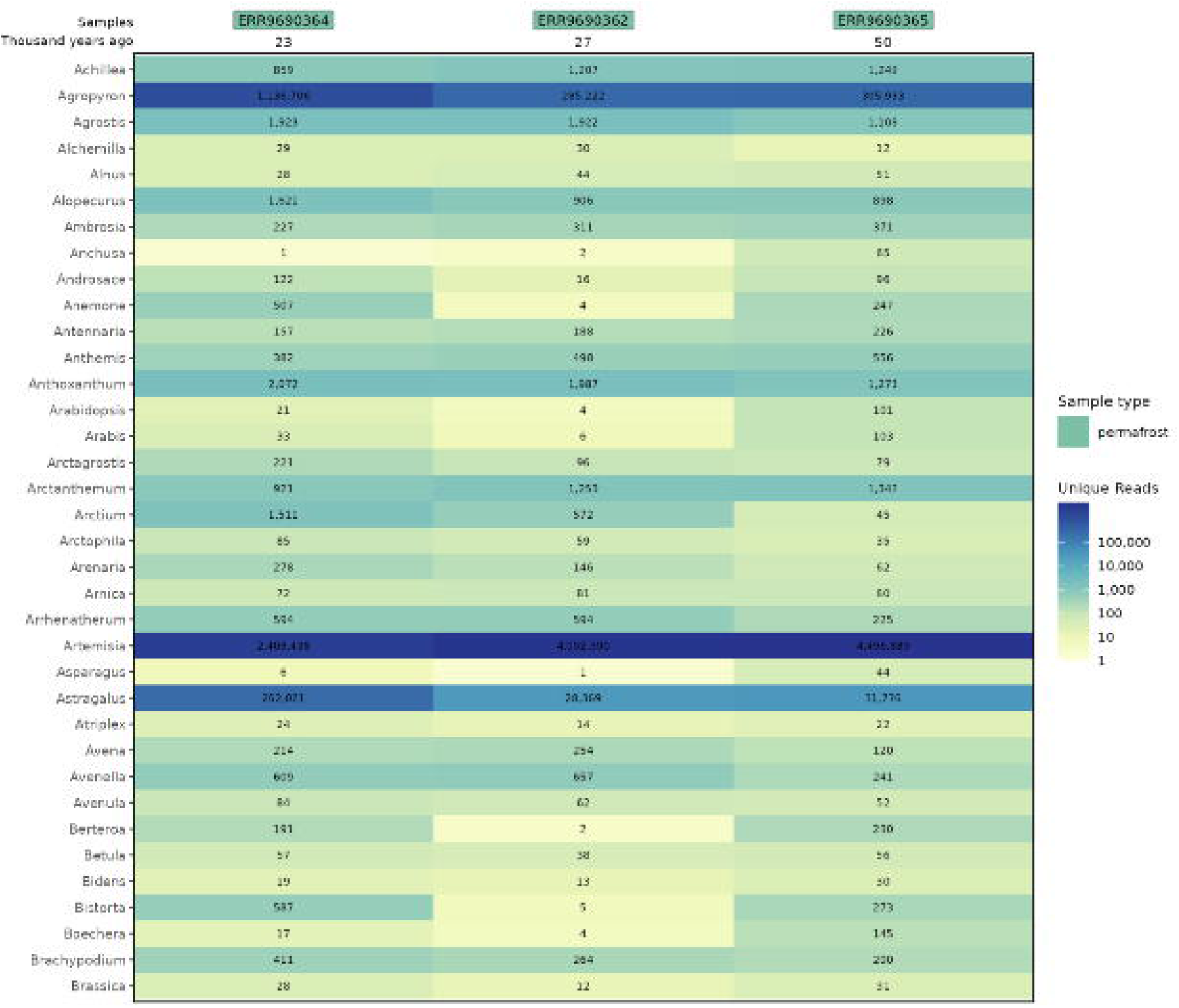
Heatmap representing the number of unique reads assigned per taxa at the genus level after the bamdam step (200 to 239)

### Mammalian composition

Different mammalian taxa were confidently detected through the metaJAM pipeline across the different samples. The Equidae family was present in the samples B17-D7 and B17-D10, with the genus Equus retrieved with a sufficient amount of reads to retrieve proper damage patterns (Supplementary Fig. 6A damage patterns). In a similar manner, the presence of Elephantidae was authenticated across the three samples (Supplementary Figs. 5 and 6B damage patterns). Most of the reads were assigned to the Elephas genus due to the presence of Elephas maximus whole genome in the custom database. Nonetheless, analysis of the MMseqs2 results suggested a correct assignment at the family level but not at the genus level (Supplementary Fig. 5), which is coherent with the detection of Mammuthus in the original analysis from Courtin et al. (2022). Only the Leporidae family was solely retrieved in metaJAM, primarily due to assignment to nuclear references. As with Viridiplantae, metaJAM also yielded more assigned reads to Mammalian taxa, which can again be explained by the use of nuclear genome references during mapping.

## Discussion

Viridiplantae and Mammalian results were successfully replicated for the three selected permafrost samples from Courtin et al. (2022) using the metaJAM workflow and demonstrate its user-friendly and functional aspects. The higher number of assigned reads found using the metaJAM pipeline is explained by the use of a different strategy than the Kraken 2 classification previously performed, specifically the addition of more comprehensive nuclear databases.

These results demonstrate the potential of the workflow, which allows the user to agilely customize the filters and databases used to best suit a given research setting. This is particularly important when computational resources are limited. The use of a database as comprehensive as possible such as the NCBI nt database, while appealing when a priori ecological information is limited or absent, currently requires more storage and CPU resources than many researchers have access to. This problem is expected to become more difficult, as the number of available reference genomes has been increasing exponentially over the last 10 years (Goldfarb et al., 2025) and shows no sign of slowing. Here, the implementation of microbial filtering and masking, along with the ngsLCA-bamdam strategy, MMseqs2 assignment, and provision of evaluation plots allows the user to rely on taxon authentication while ruling out and investigating potential false assignments. These steps allow users to ultimately receive a confident overview of the reconstructed eukaryotic taxa assemblages in metagenomic datasets while limiting resource use.

In particular, the significantly decreased computation time of MMseqs2 compared to BLAST (Kallenborn et al., 2025; Steinegger and Söding, 2017) allows users to quickly query a restrained set of reads assigned to taxa of interest to the most recent nt-database to confirm or improve taxonomic assignments. This limits the need to rely on generalized, massive reference datasets during the initial mapping step, as assignments based on smaller, but better curated databases can be more easily assessed.

Furthermore, the flexibility in how metaJAM can be configured allows it to be used not only for sedimentary ancient DNA, but it could also be used for more single-taxa-oriented research, such as the detection of the provenance of unidentified bones or bulk bones, such as those found in archaeological contexts (e.g. Seersholm et al., 2022; Xing et al., 2025). The combination of reduced computational resource use, flexibility in workflow design, and ease of set-up through Nextflow are therefore the major strengths of metaJAM.

Among the different existing options that arise when choosing a bioinformatic pipeline to process ancient DNA metagenomic samples, the Holi pipeline (Pedersen et al., 2016; Vogel et al., 2026) is a very effective one that is the more similar in term of mapping strategy to metaJAM, while nf-core/eager (Yates et al., 2021) is closest in spirit to metaJAM through its Nextflow implementation and its flexibility in tool selection. Nevertheless, the introduction in metaJAM of reference genomes microbial masking, iterative or parallele competitive mapping, the integration of up-to-date metagenomic filtering/plotting tools, and additional validation with blastn-like software complemented by diagnostic plots, makes it an attractive option for a wide range of researchers seeking to process eukaryotic analysis of their metagenomic ancient samples with relative ease.

## Acknowledgements

We thank Peter D. Heintzman and Anna Linderholm for their advice and suggestions regarding the workflow. We also thank the members of the Center for Palaeogenetics in Stockholm for testing the pipeline and providing valuable feedback, notably Flore Wijnands, Lauren Clark, Inda Brinkmann and Scott Cocker.

## Author Contributions

E.J, J.A and N.M conceived the ideas and designed methodology; E.J, J.A, C.J, B.G and N.M created the workflow; E.J, C.J.,J.A and N.M analysed the data and N.M. led the writing of the manuscript. All authors contributed critically to the writing of the drafts and gave final approval for publication.

## Data Availability

The workflow is available on https://github.com/NathanACO/metaJAM/tree/metaJAM-nf.

---

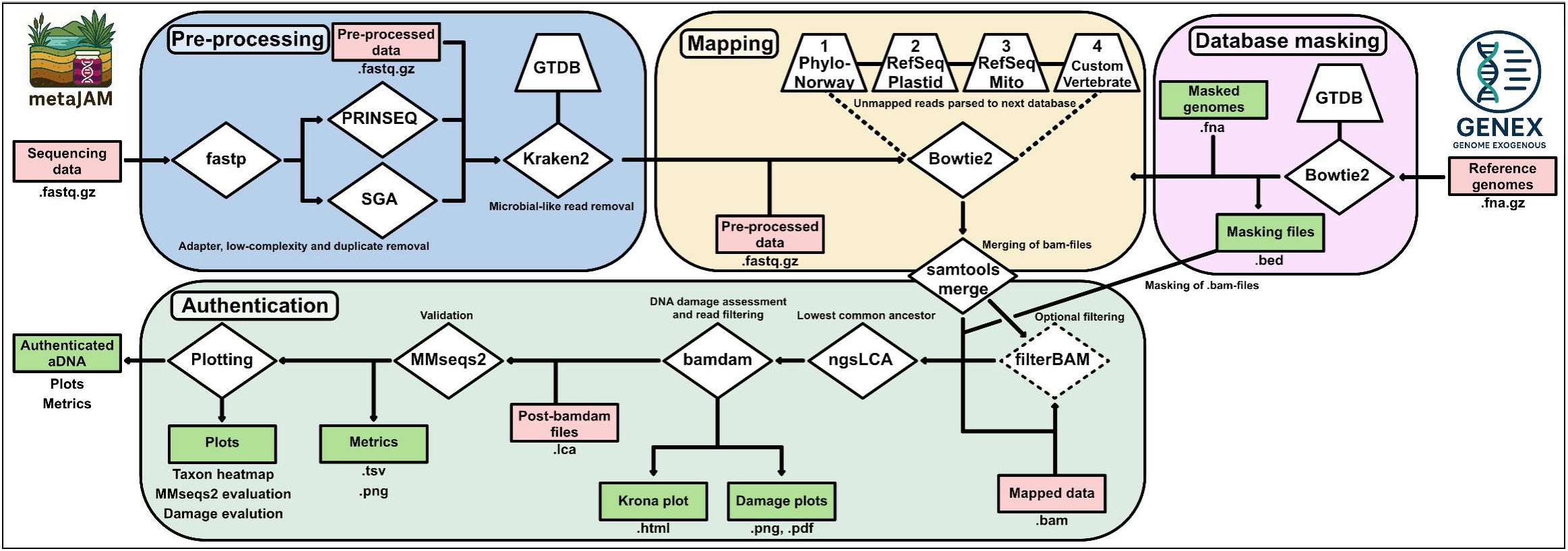

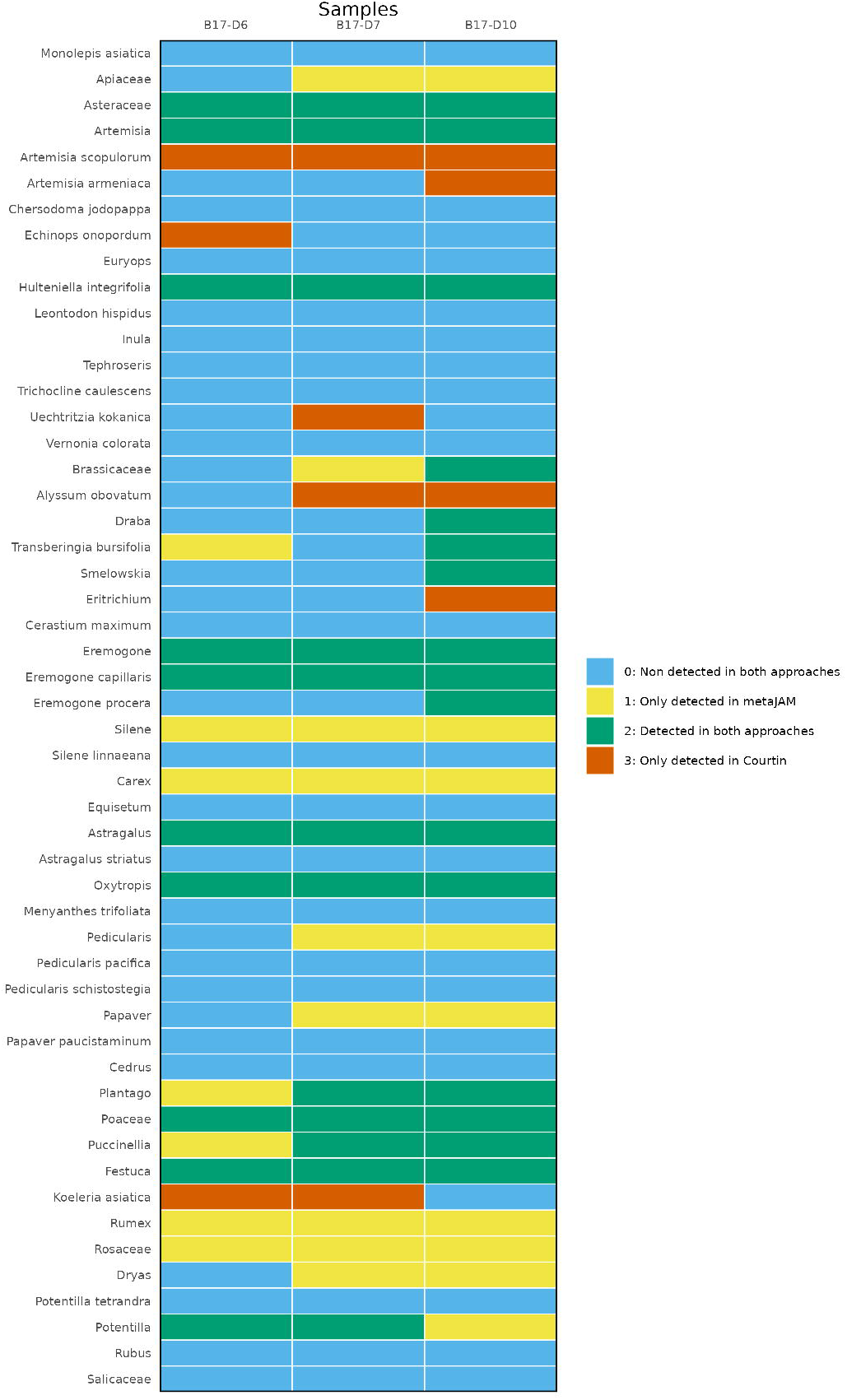

