## Supplementary figures and images for "metaJAM: a Nextflow integrated metagenomic workflow for sedimentary ancient DNA"

### Supplementary Figure 1

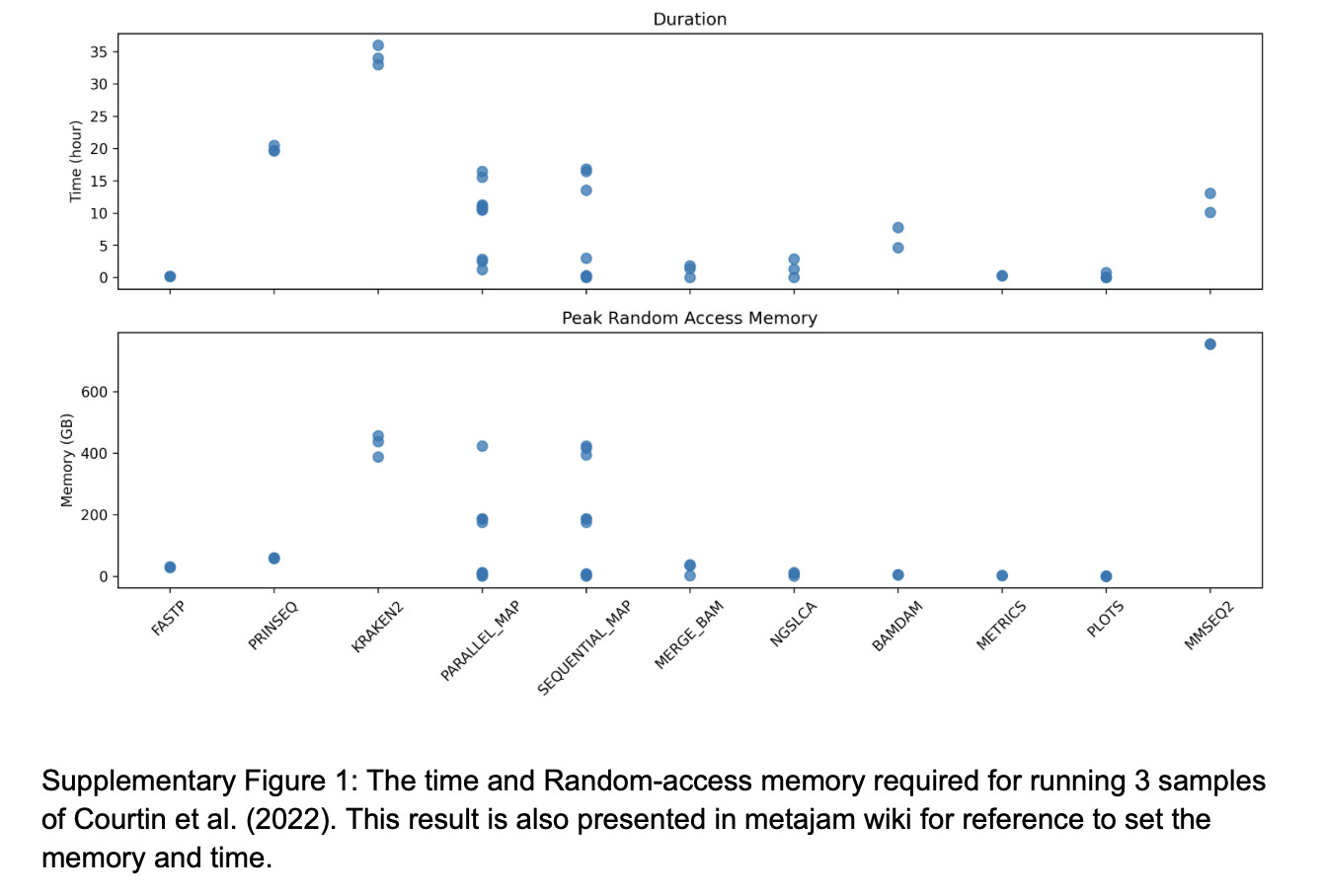

### Supplementary Figure 2

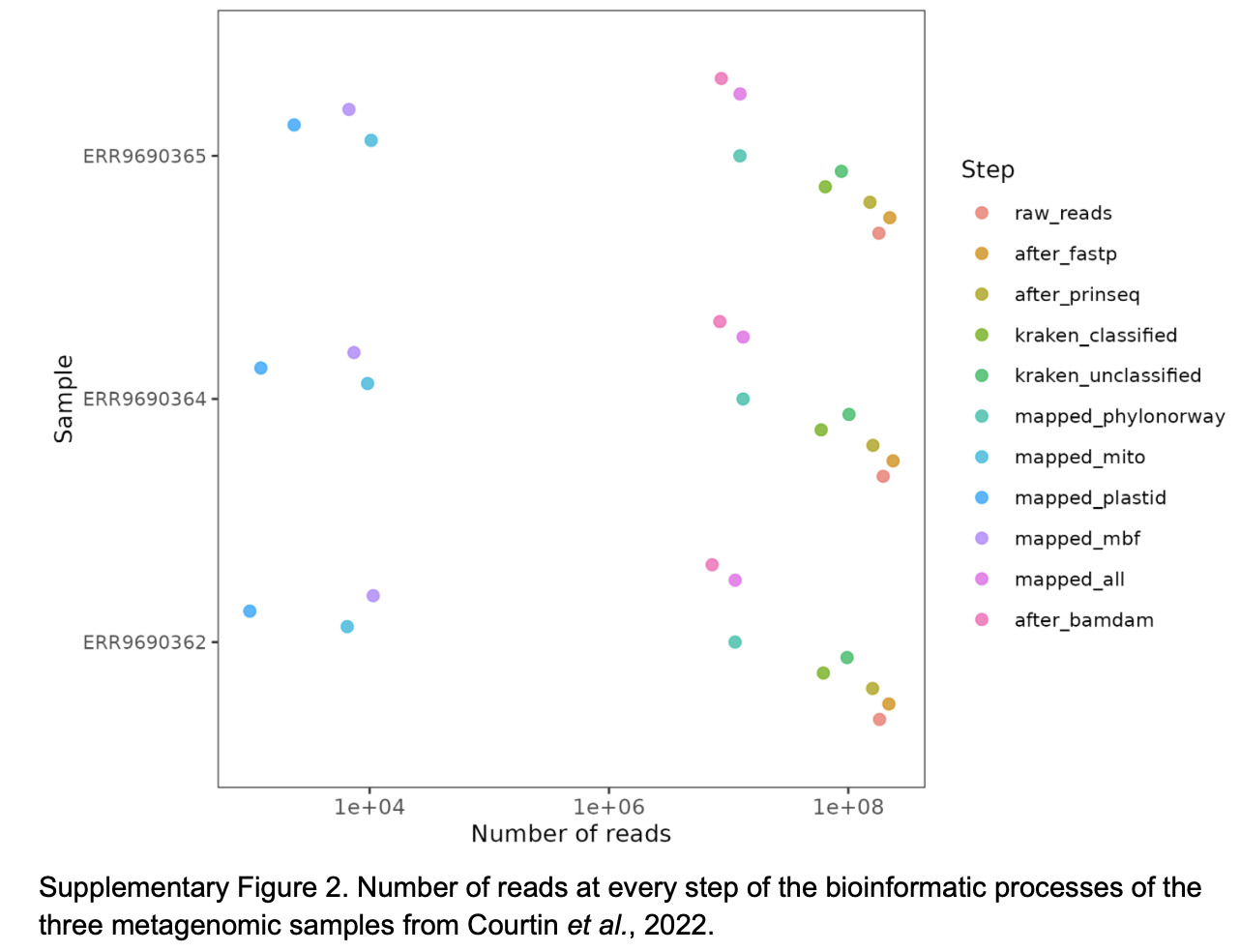

### Supplementary Figure 3.A

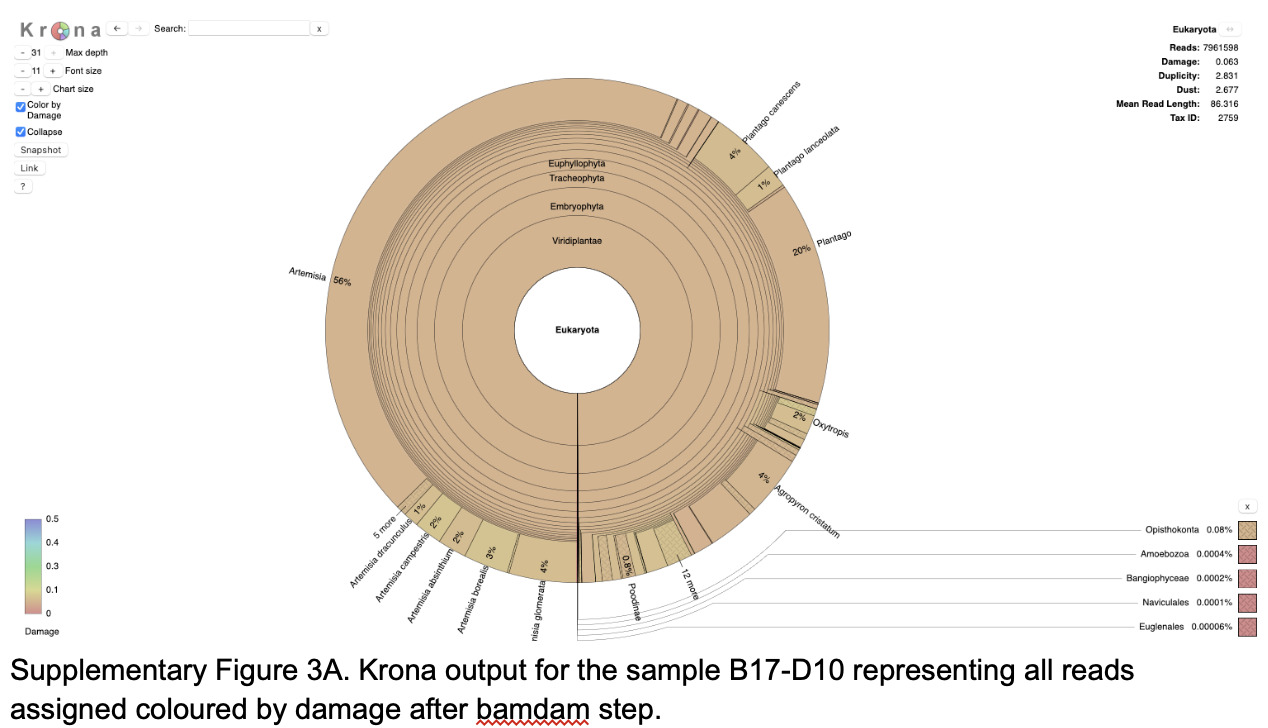

### Supplementary Figure 3.B

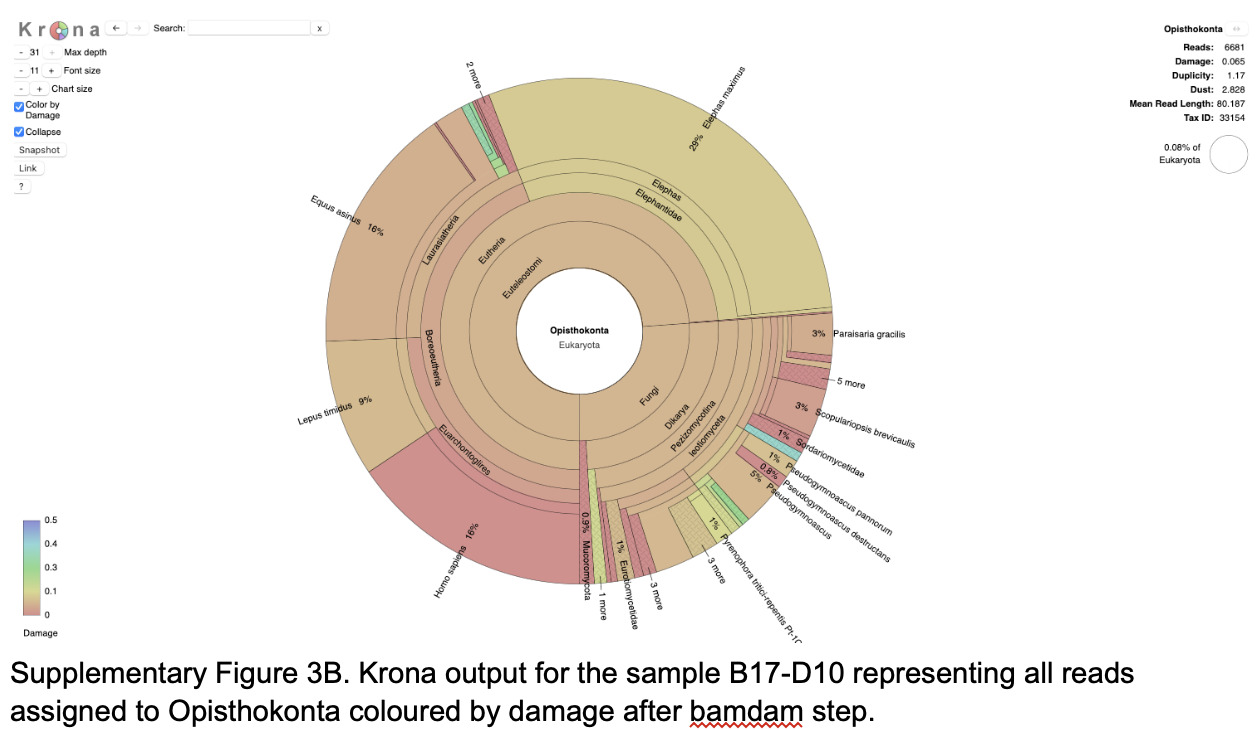

### Supplementary Figure 4.A

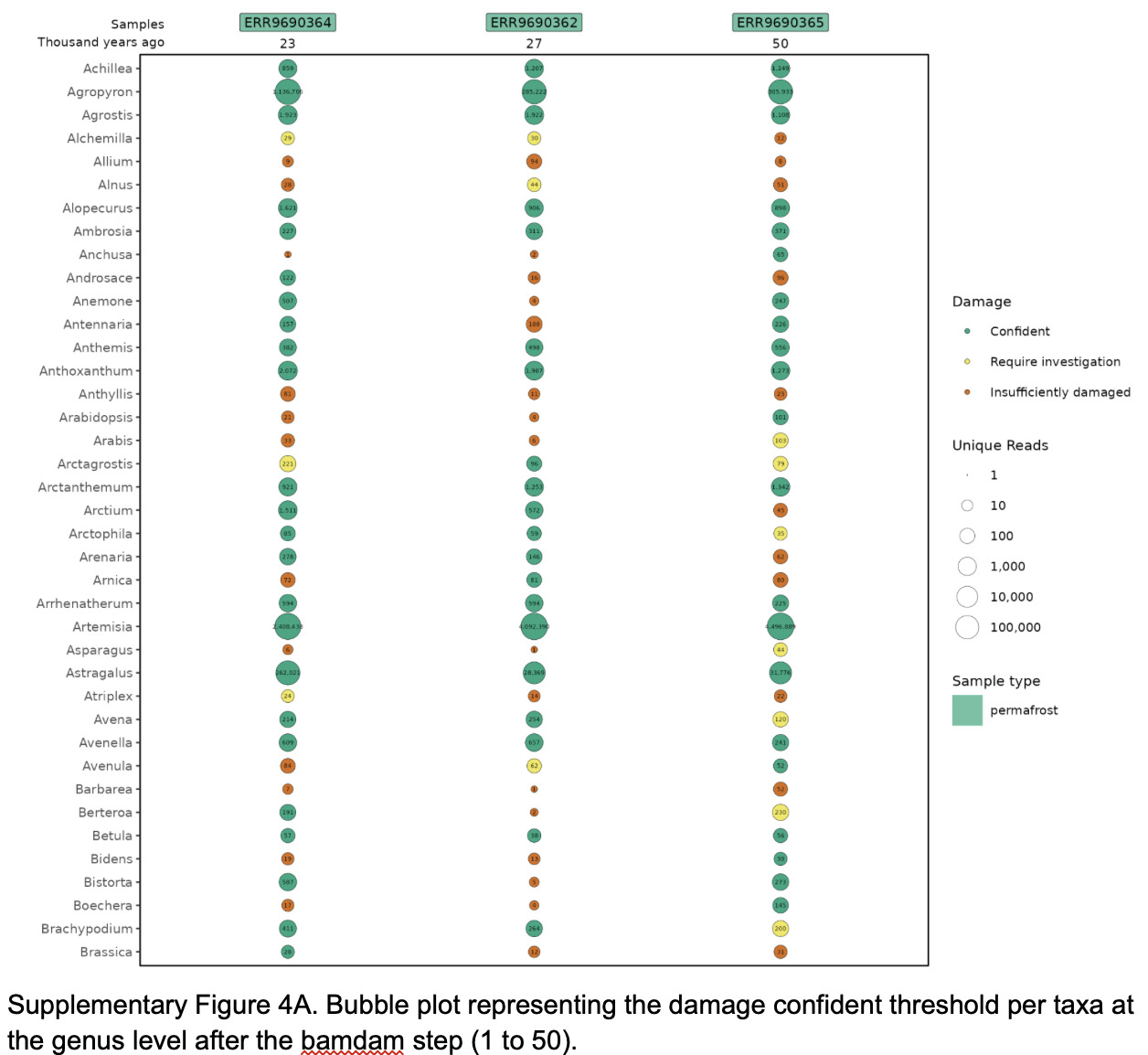

### Supplementary Figure 4.B

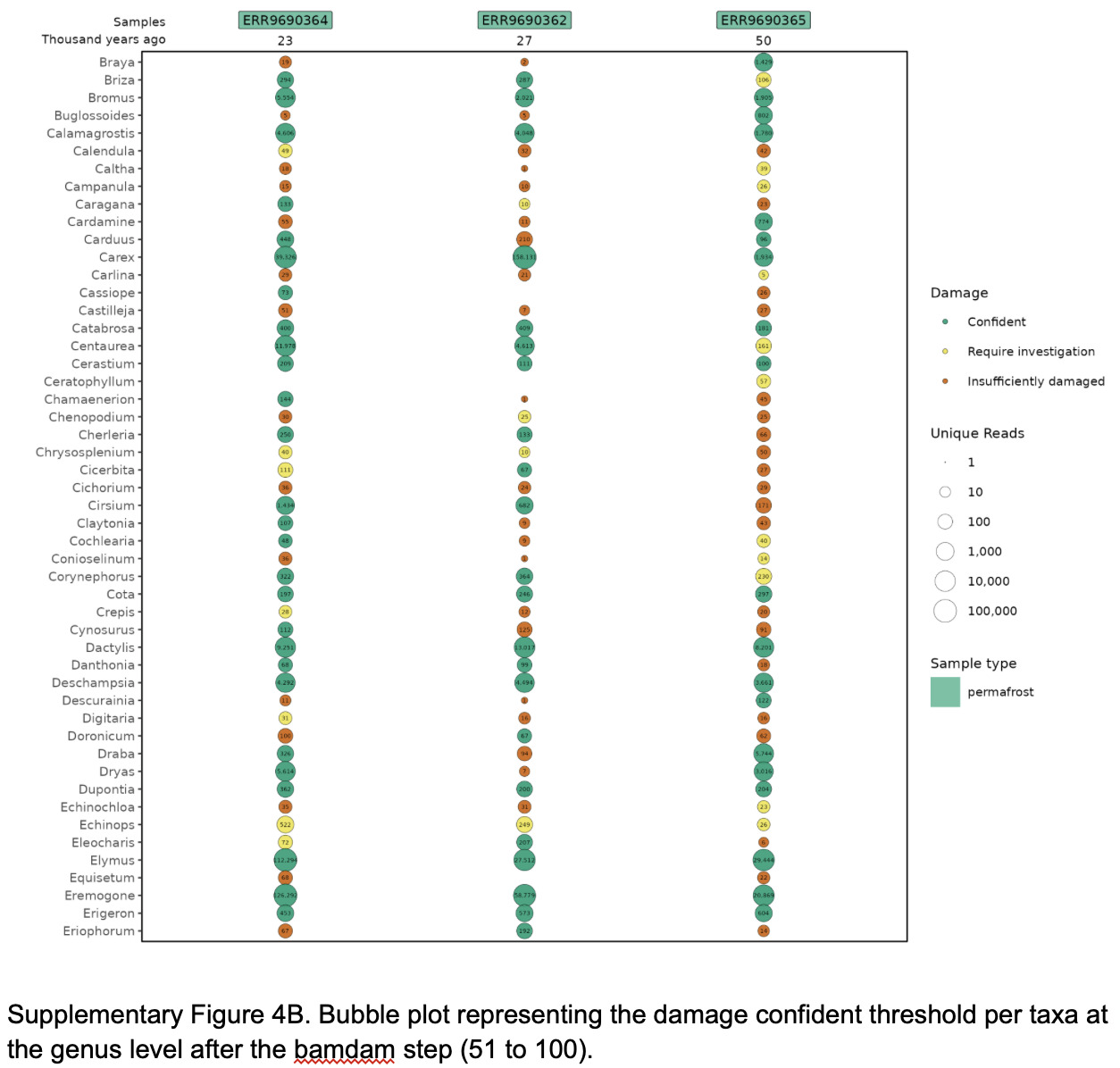

### Supplementary Figure 4.C

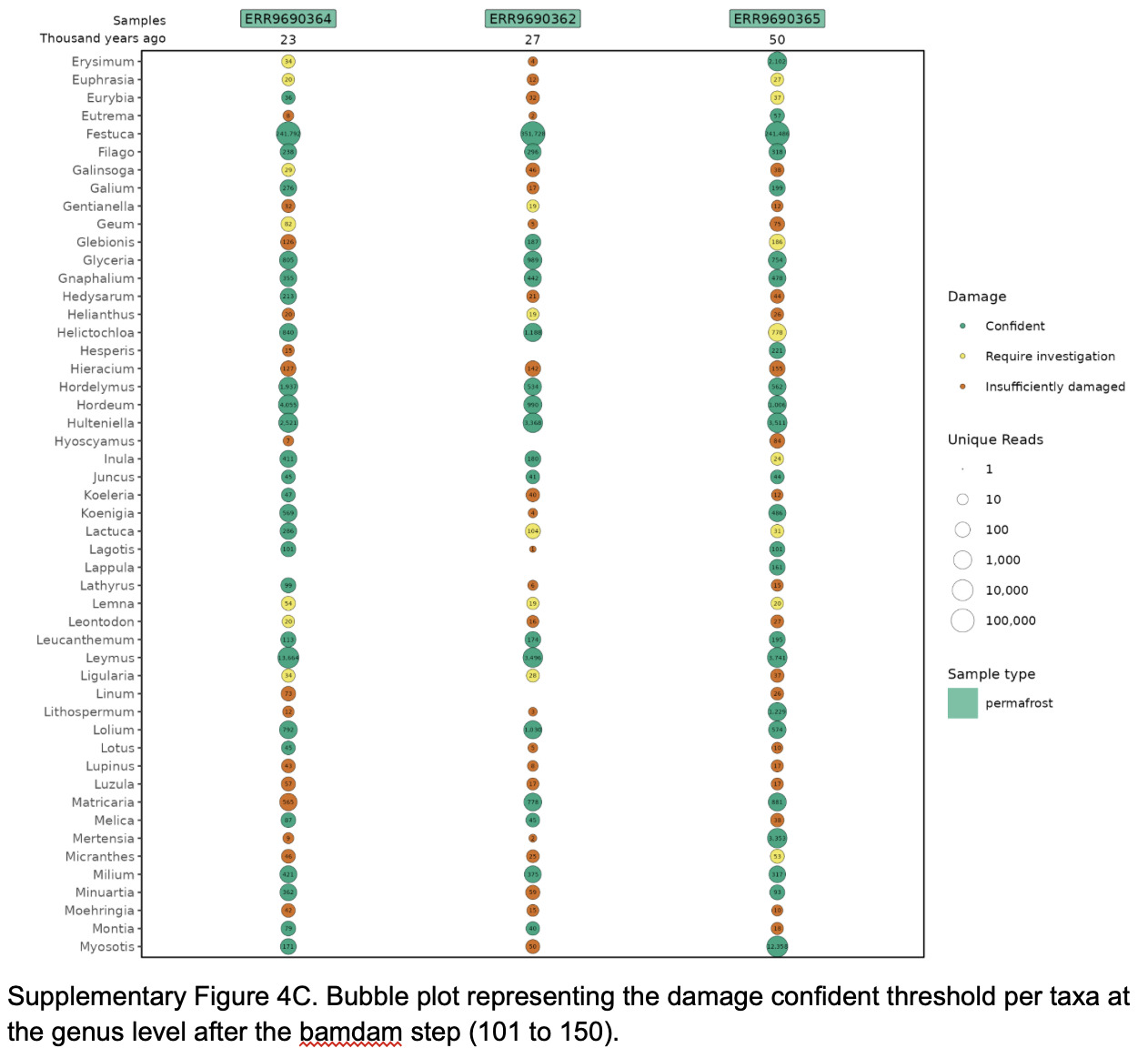

### Supplementary Figure 4.D

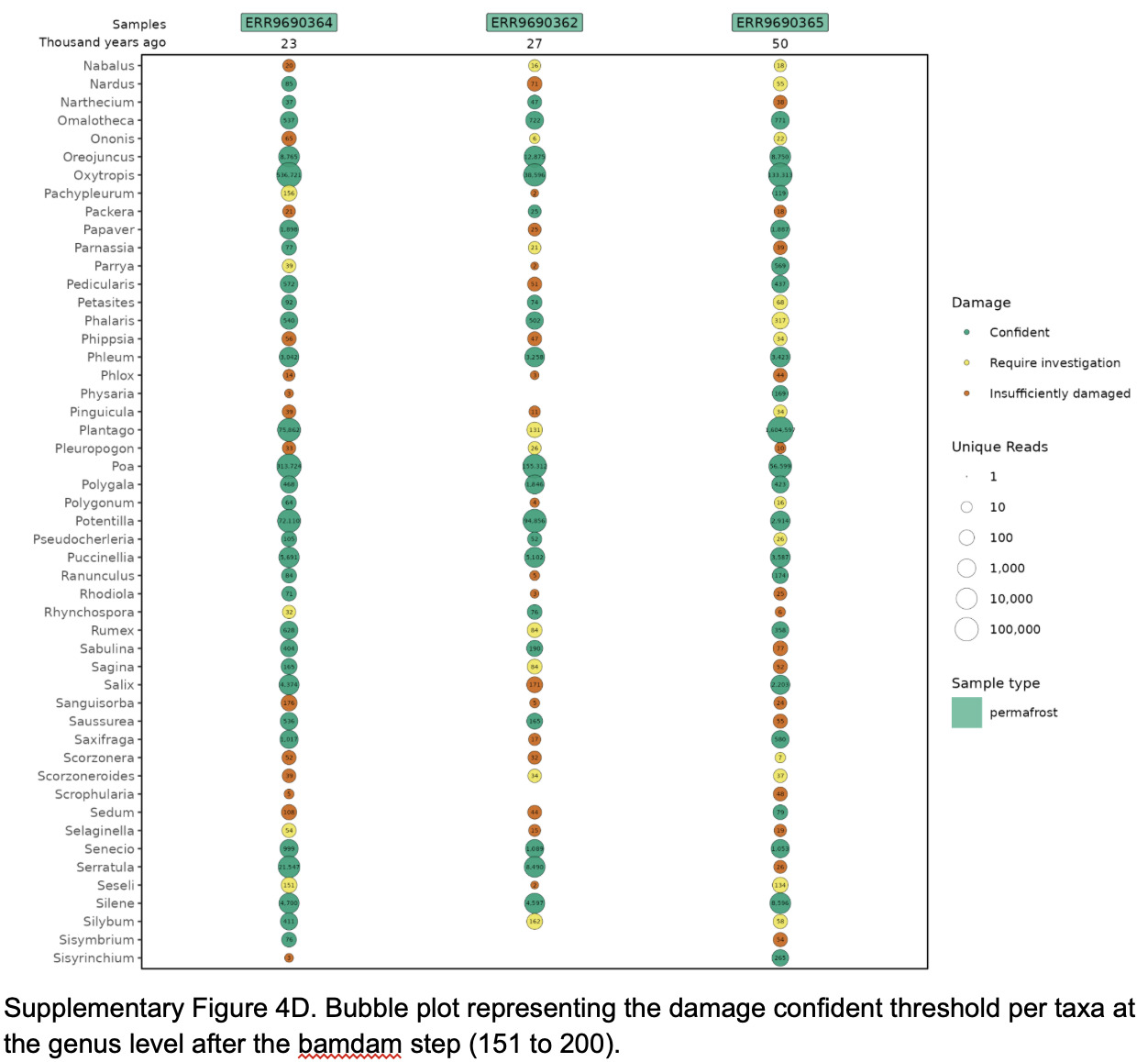

### Supplementary Figure 4.E

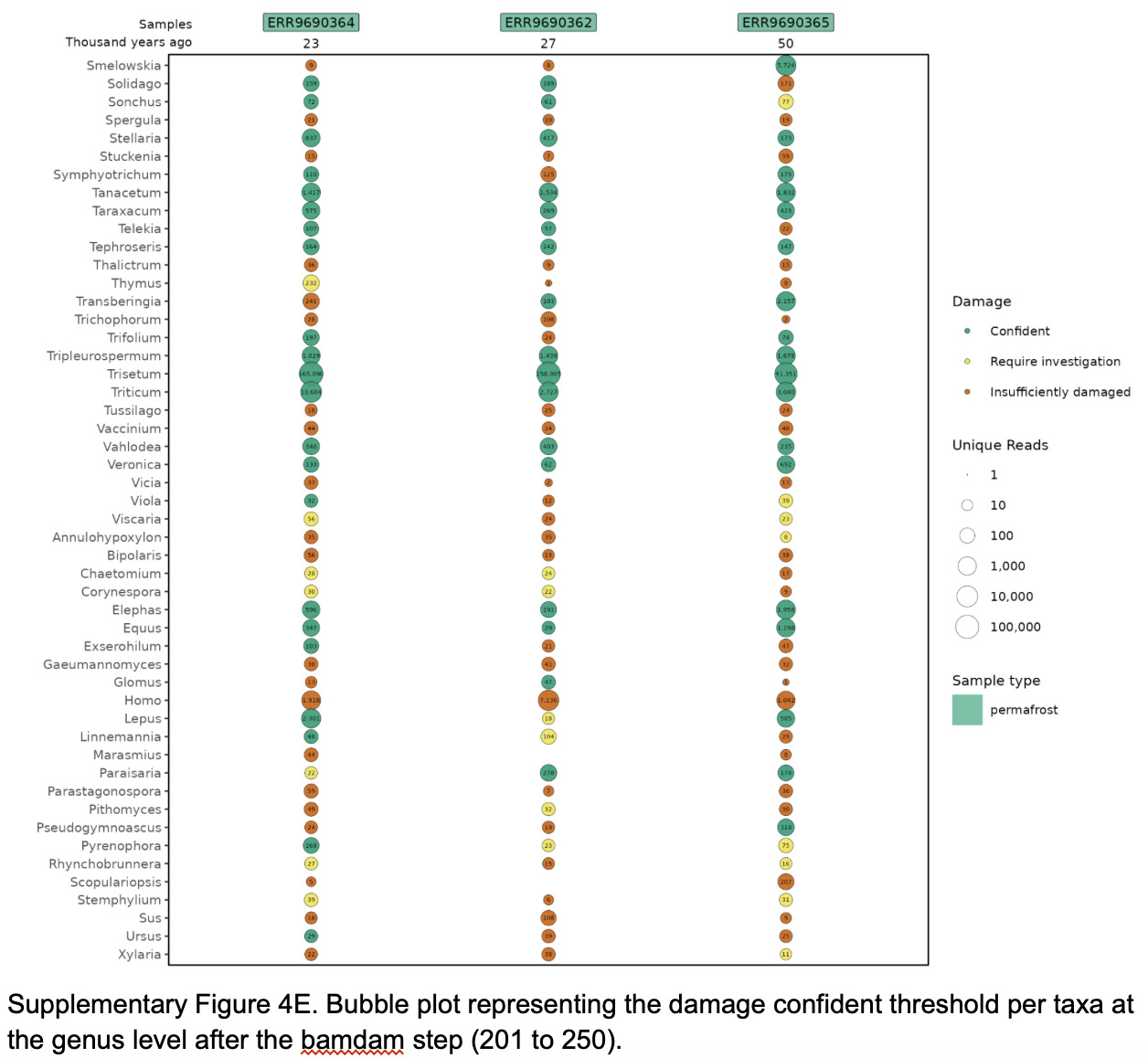

### Supplementary Figure 4.F

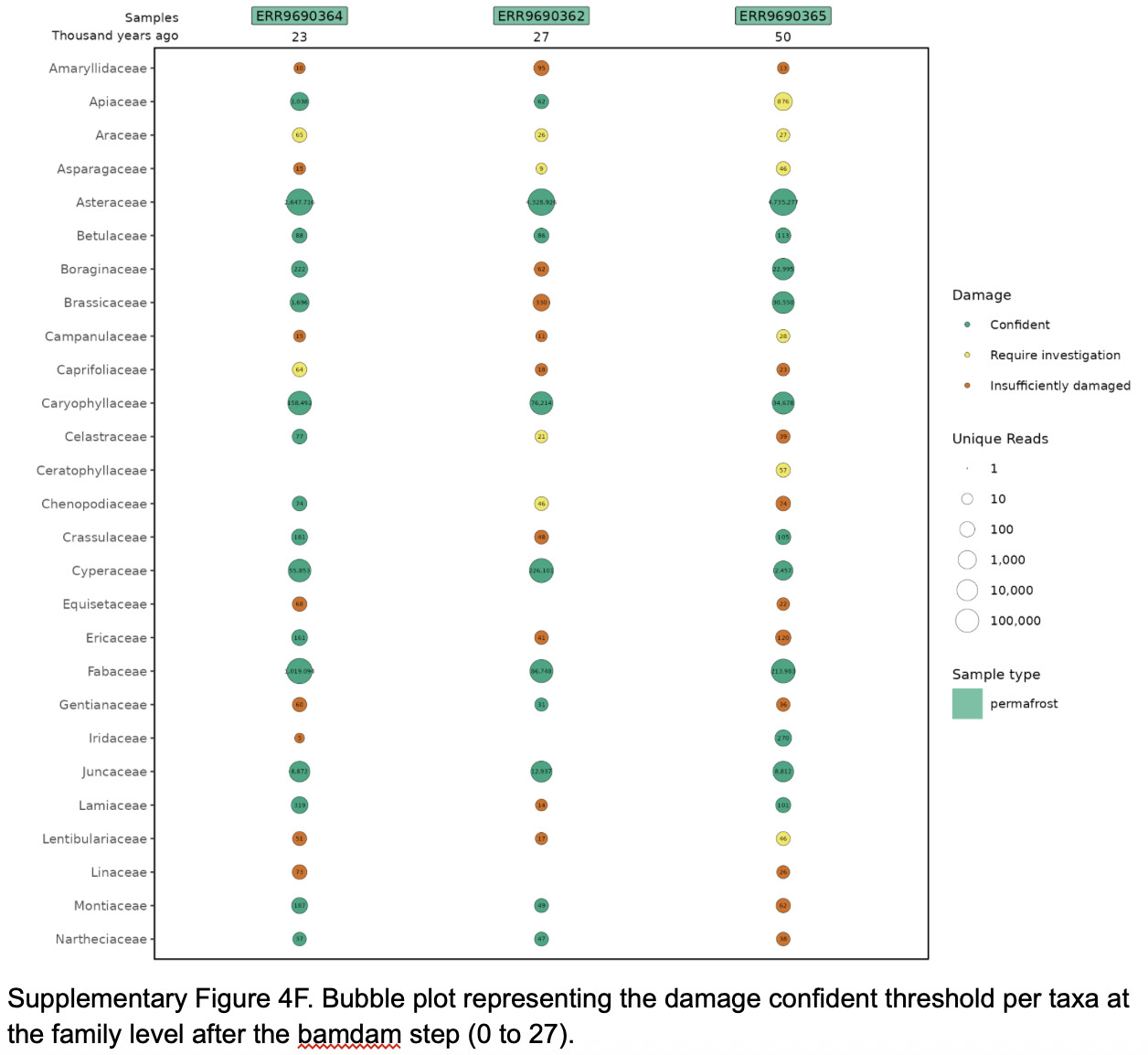

### Supplementary Figure 4.G

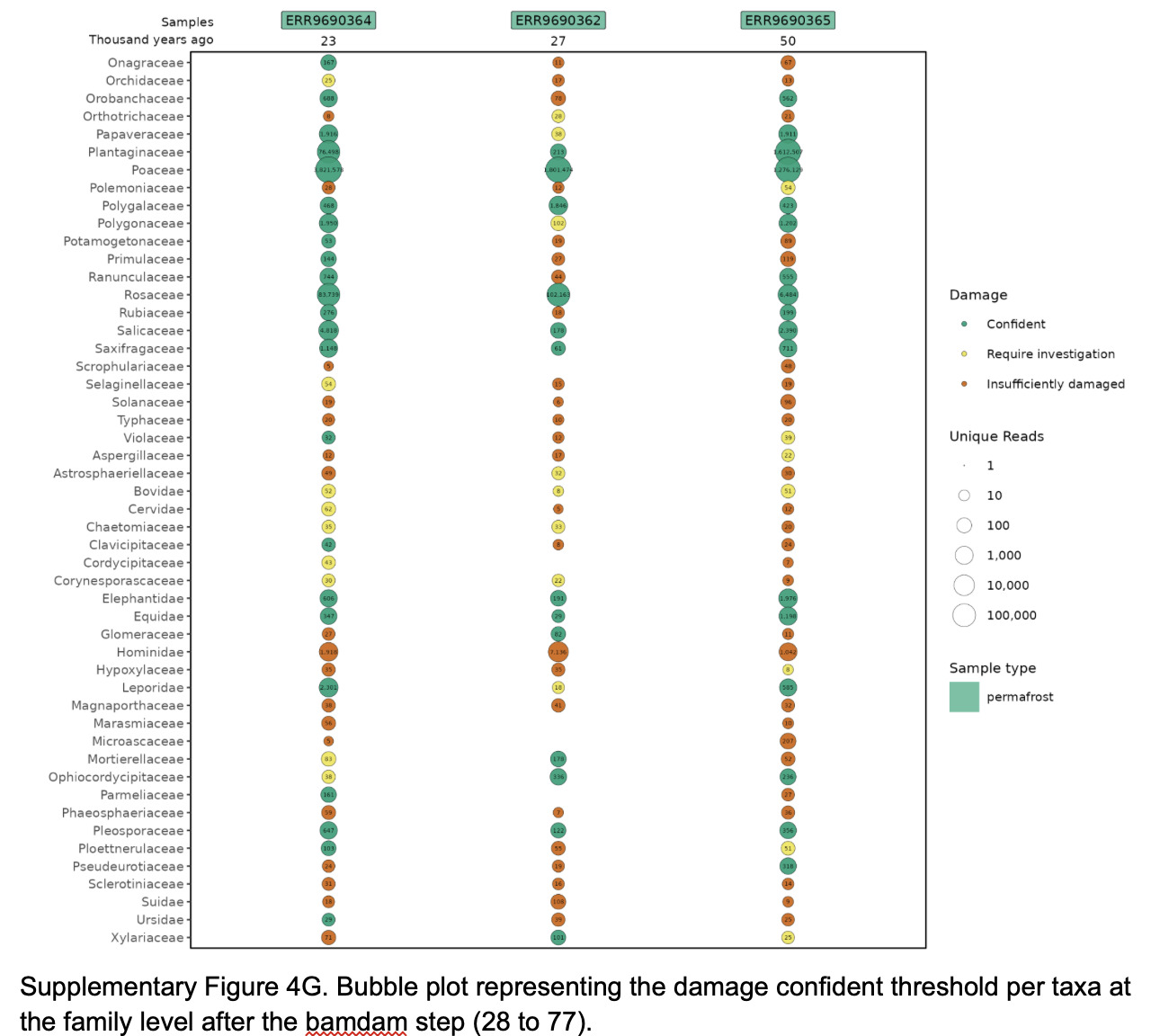

### Supplementary Figure 5.A

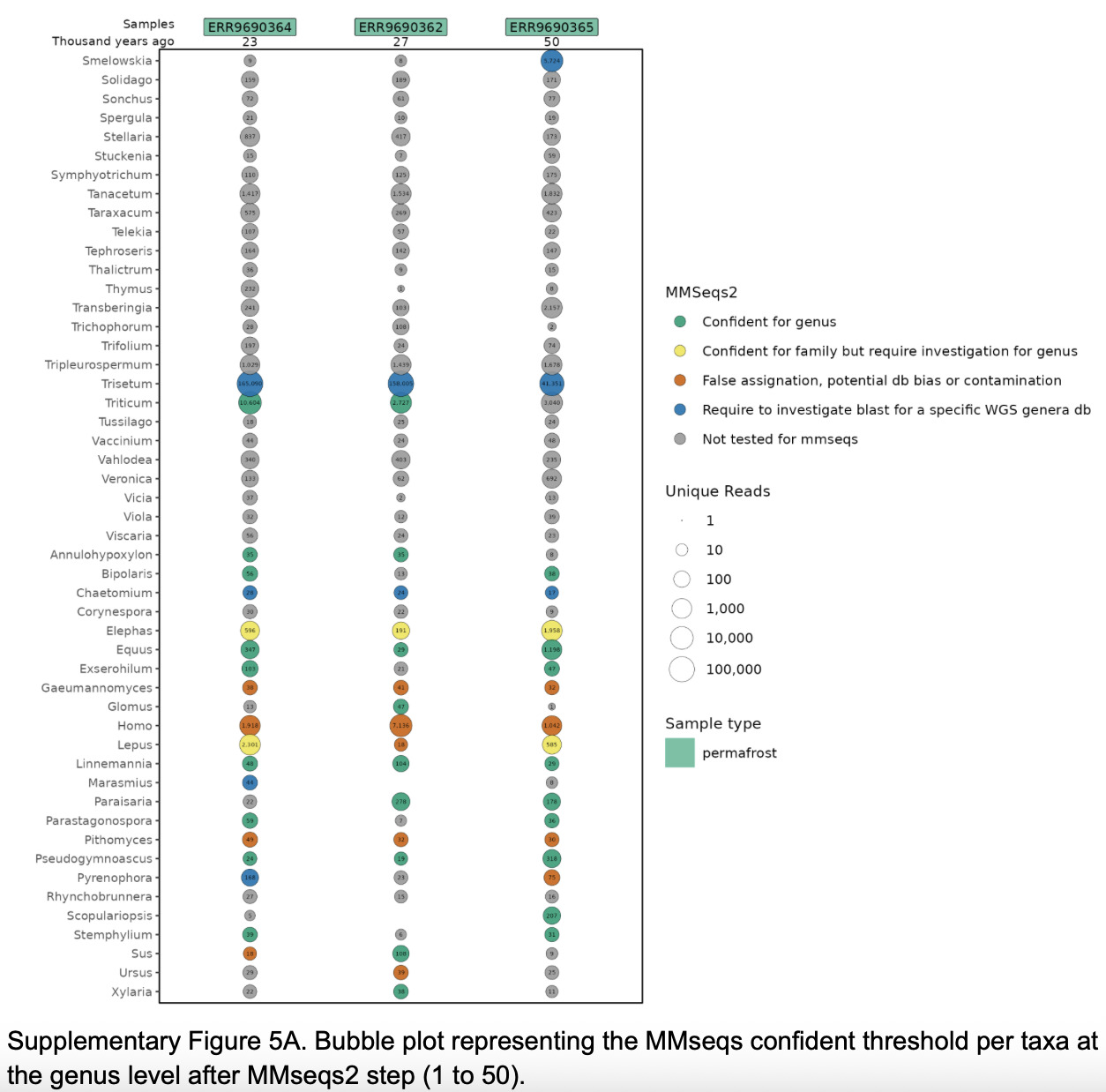

### Supplementary Figure 5.B

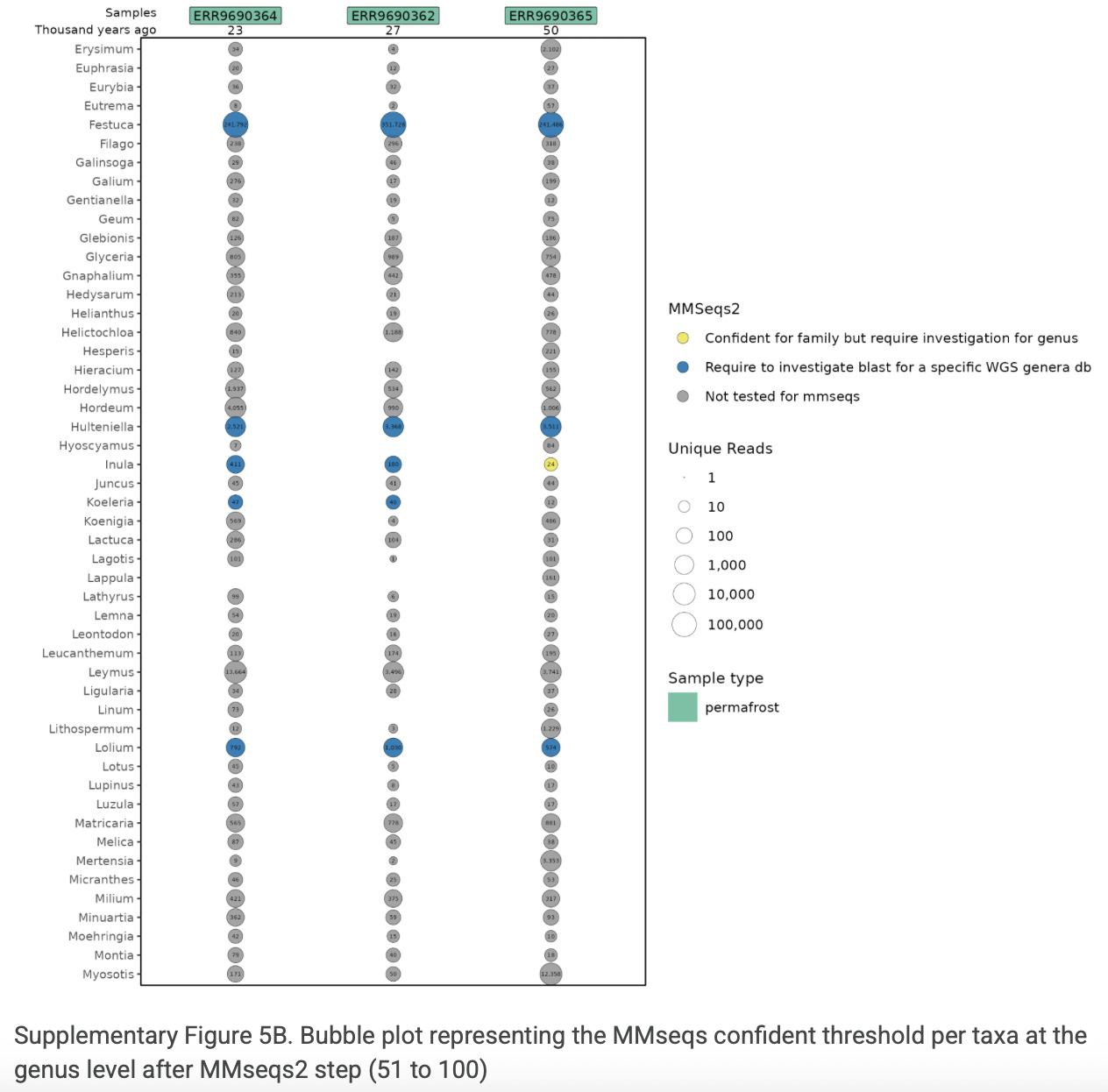

### Supplementary Figure 5.C

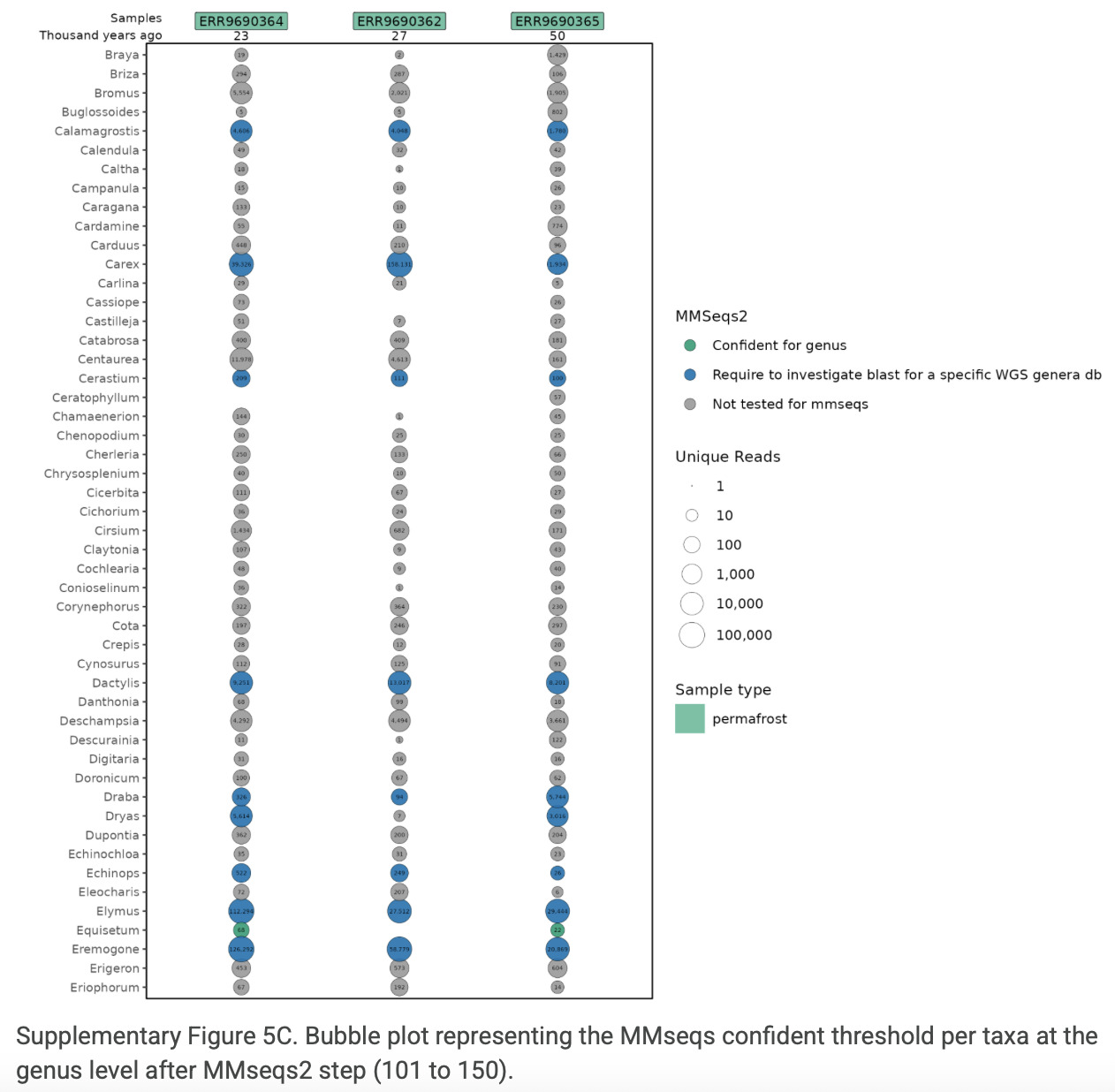

### Supplementary Figure 5.D

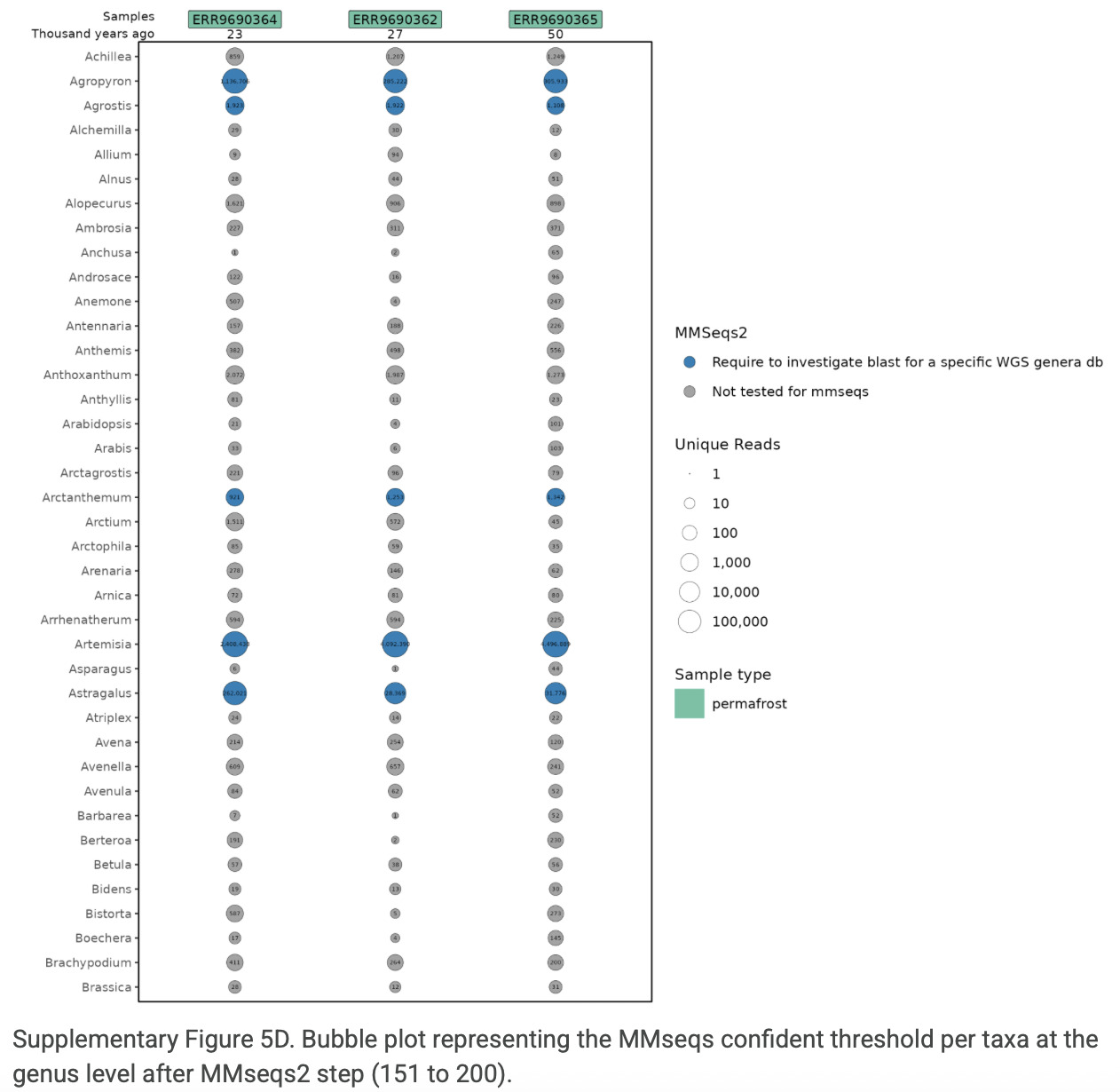

### Supplementary Figure 5.E

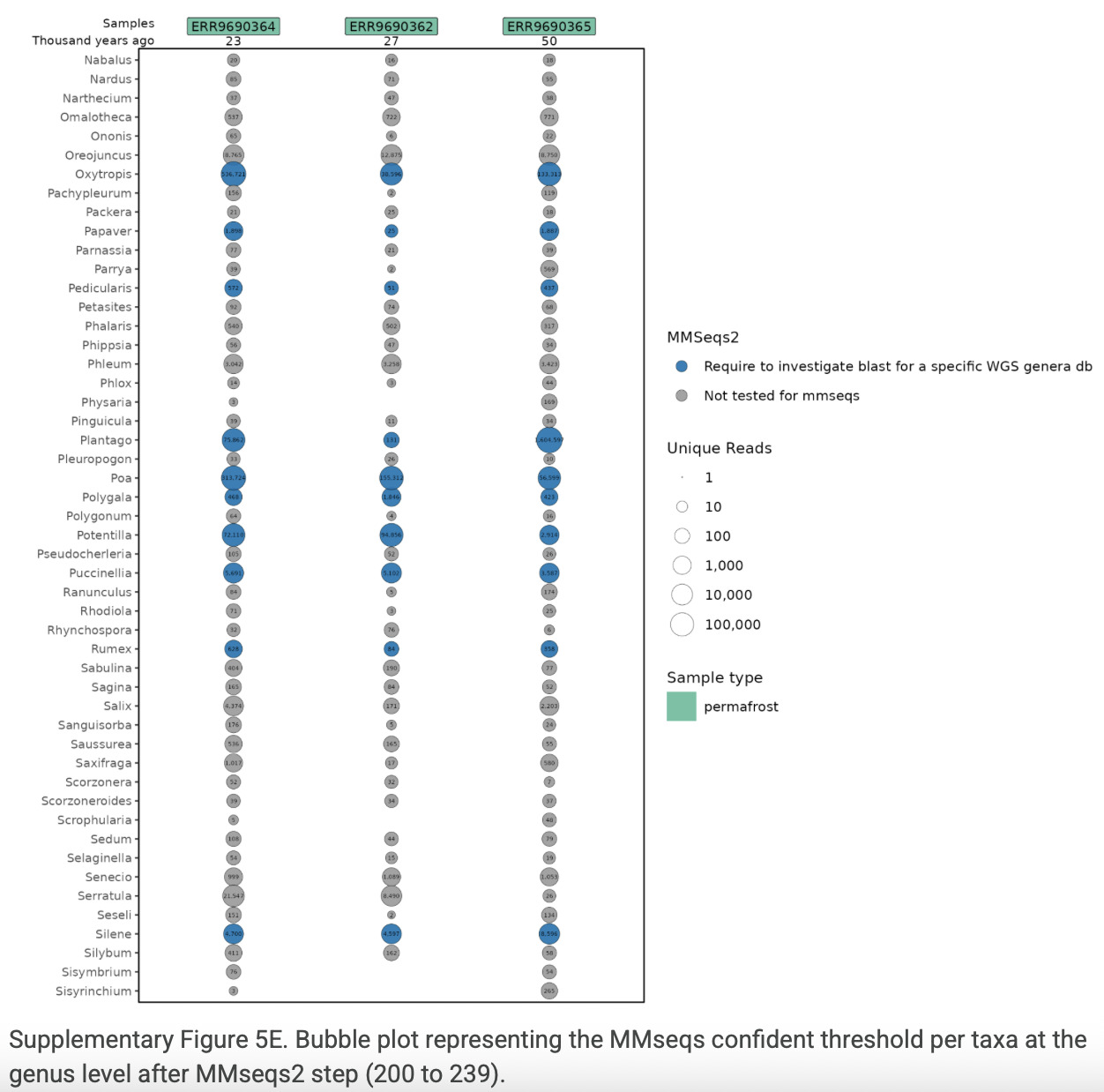

### Supplementary Figure 6.A

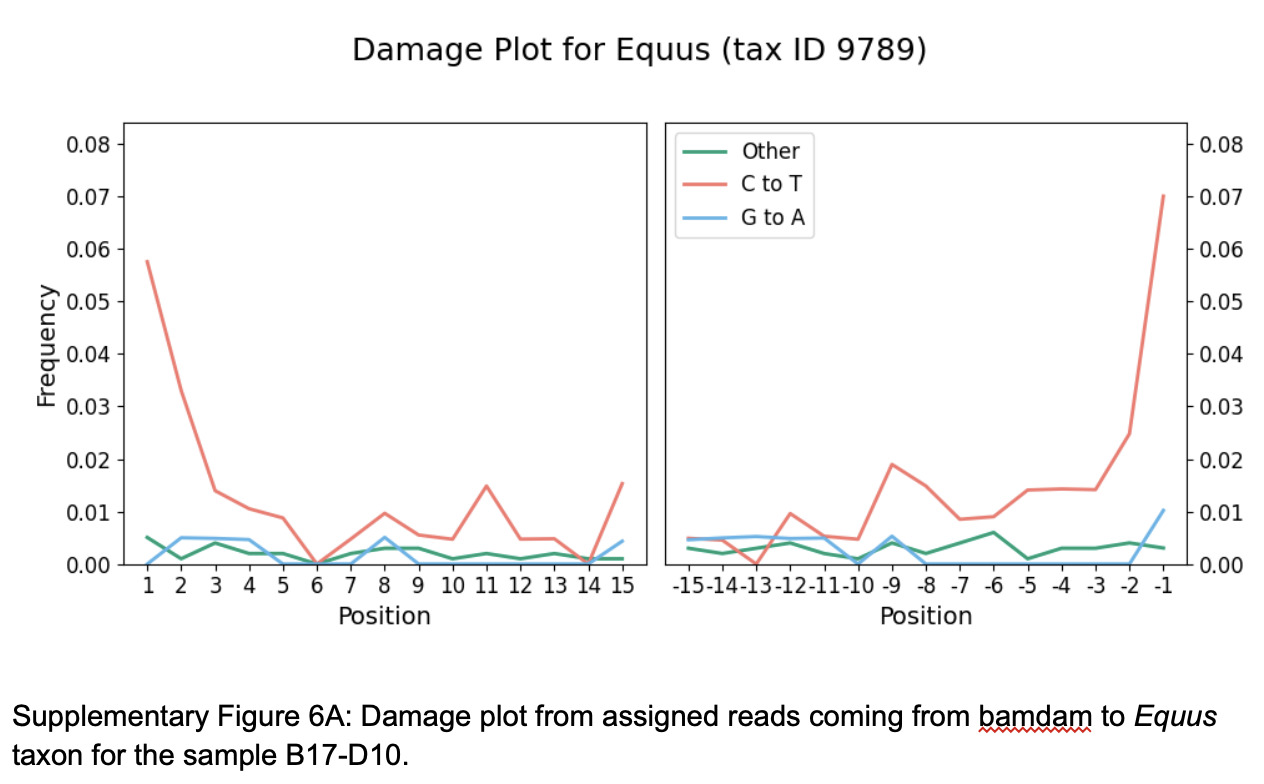

### Supplementary Figure 6.B

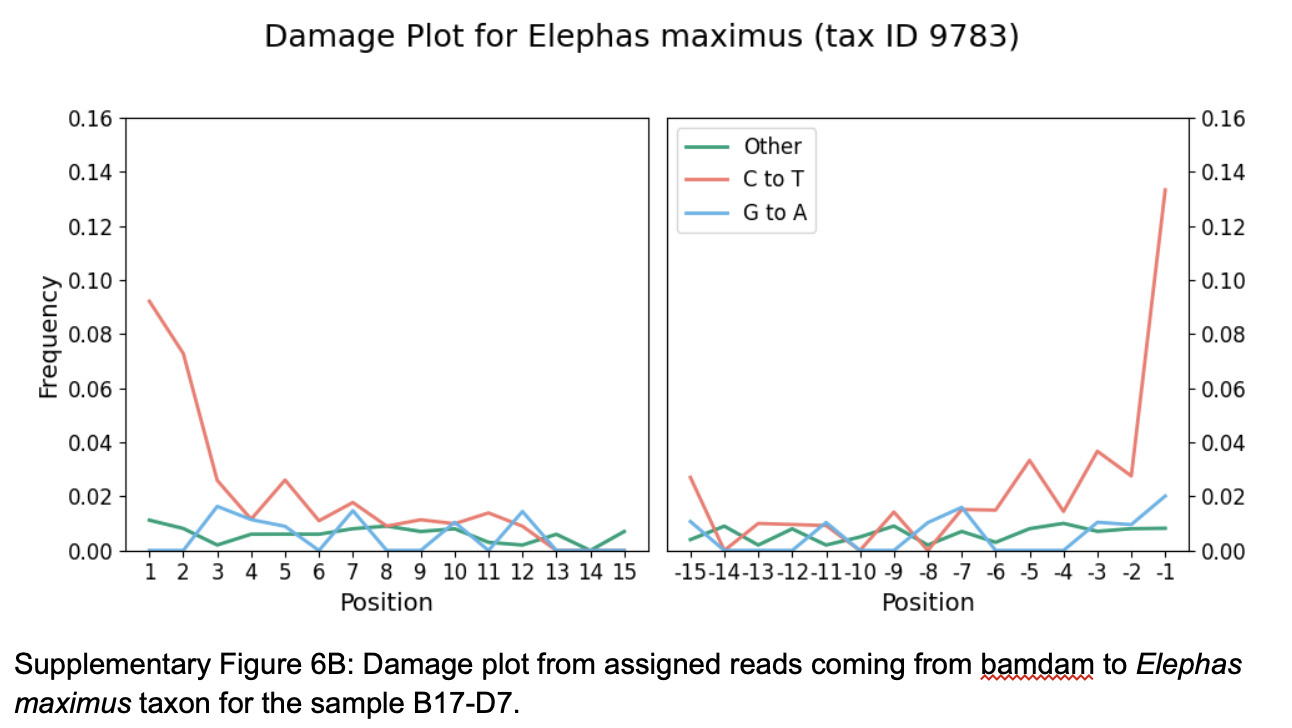

### Supplementary Figure 6.C

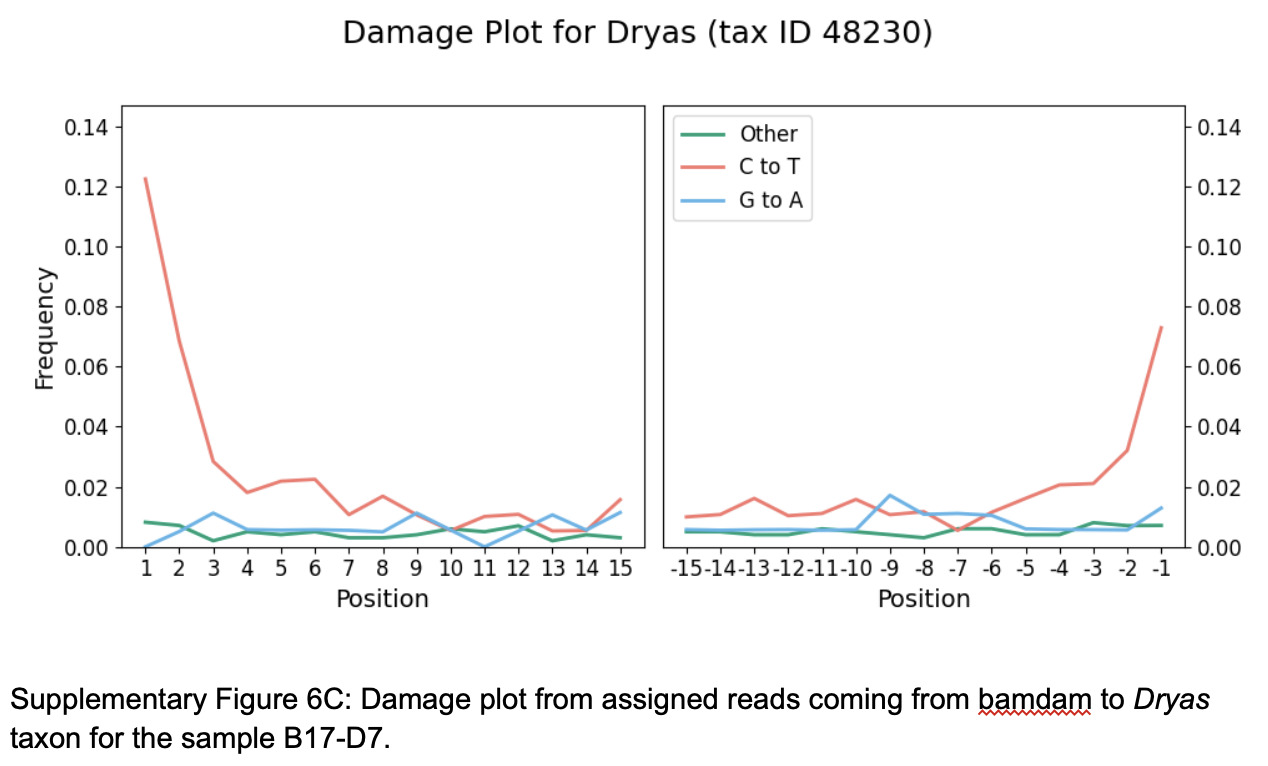

### Supplementary Figure 6.D

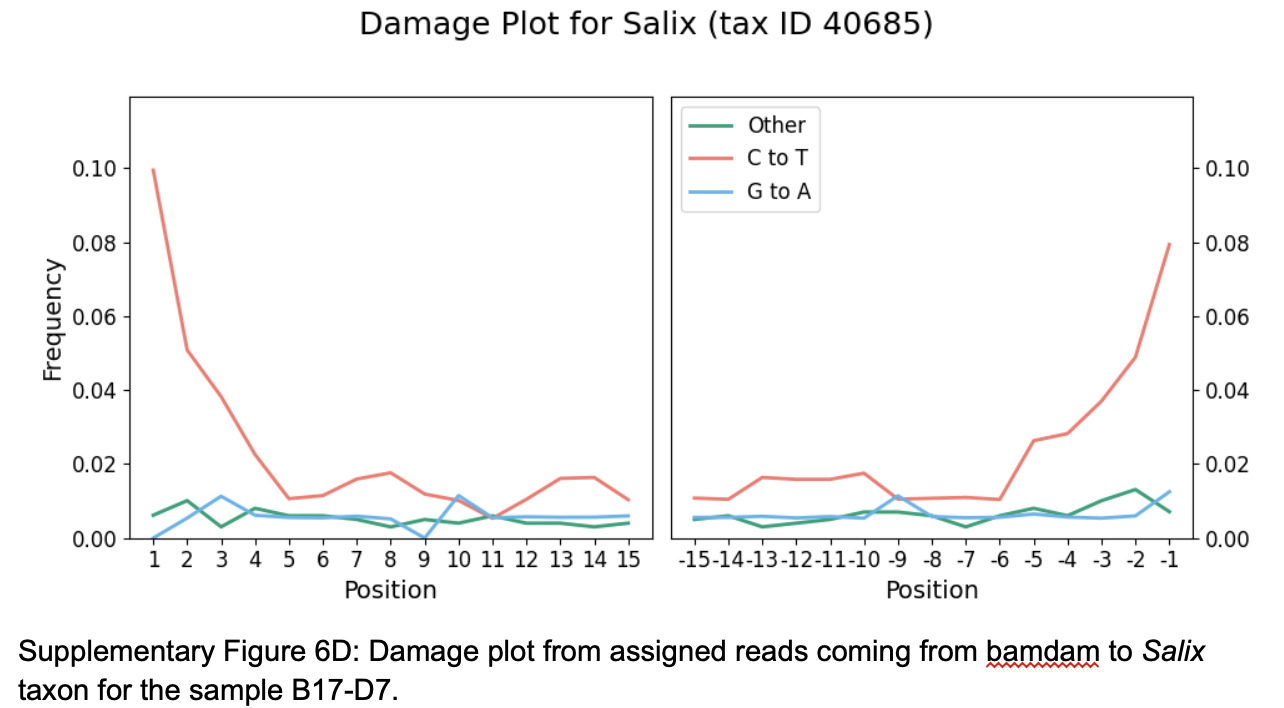

### Supplementary Figure 6.E

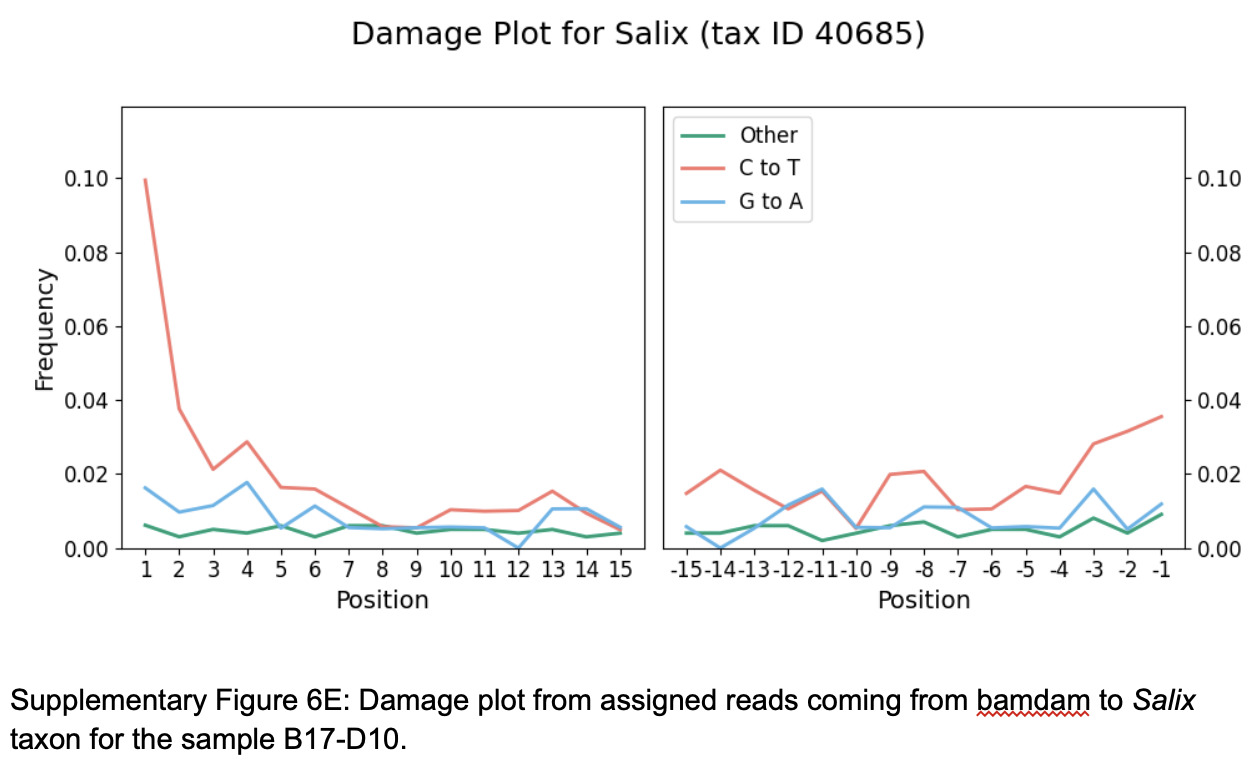
